# Cell Lineage and Communication Network Inference via Optimization for Single-cell Transcriptomics

**DOI:** 10.1101/168922

**Authors:** Shuxiong Wang, Matthew Karikomi, Adam L. MacLean, Qing Nie

**Author notes:** To whom correspondence should be addressed. Correspondence may also be addressed to Qing Nie.

## Abstract

The use of single-cell transcriptomics has become a major approach to delineate cell subpopulations and the transitions between them. While various computational tools using different mathematical methods have been developed to infer clusters, marker genes, and cell lineage, none yet integrate these within a mathematical framework to perform multiple tasks coherently. Such coherence is critical for the inference of cell-cell communication, a major remaining challenge. Here we present similarity matrix-based optimization for single-cell data analysis (SoptSC), in which unsupervised clustering, pseudotemporal ordering, lineage inference, and marker gene identification are inferred via a structured cell-to-cell similarity matrix. SoptSC then predicts cell-cell communication networks, enabling reconstruction of complex cell lineages that include feedback or feedforward interactions. Application of SoptSC to early embryonic development, epidermal regeneration, and hematopoiesis demonstrates robust identification of subpopulations, lineage relationships, and pseudotime, and prediction of pathway-specific cell communication patterns regulating processes of development and differentiation.

## 1 Introduction

Our ability to measure the transcriptional state of a cell — and thus interrogate cell states and fates (1, 2) — has advanced dramatically in recent years (3) due in part to high-throughput single-cell RNA sequencing (scRNA-seq) (4). This shift, permitting delineation of different sources of heterogeneity (5, 6), requires appropriate dimension reduction techniques, cell clustering, pseudotemporal ordering of cells, and lineage inference.

Many clustering methods have been used to identify cell subpopulations via some combination of dimensionality reduction and learning of cell-to-cell similarity measures that best capture relationships between cells from their high dimensional gene profiles. Seurat and CIDR, for example, first embed single-cell gene expression data into low dimensional space by principal components analysis (PCA), and then cluster cells using a smart local moving algorithm, or hierarchical clustering, respectively (7, 8). SIMLR learns a cell-cell similarity matrix by fitting the data with multiple kernels, before using spectral clustering to identify cell subpopulations (9). An alternative recent method, SC3, constructs a cell-cell consensus matrix by combining multiple clustering solutions, and then performs hierarchical clustering with complete agglomeration on this consensus matrix (10). Cell subpopulations can also be identified using machine learning approaches (11, 12) or by analyzing cell-specific gene regulatory networks (13). The number of subpopulations is usually required as input, but can also be determined by statistical approaches (10) or via the eigengap of the cell-cell similarity matrix (9). Unsupervised prediction of the number of cell subpopulations from data remains challenging.

Marker genes - the genes that best discriminate between cell subpopulations - can be estimated by differential gene expression analysis between pairs of subpopulations (14). For example, SIMLR uses the Laplacian score to infer marker genes for each cell subpopulations (9). SC3 infers marker genes by creating a binary classifier for each gene based on mean cluster-expression values and ranking genes within clusters by highest mean expression values (10). Currently, most methods for marker gene identification (e.g. (7, 10)) are carried out *after* clustering and identification of the cell subpopulations, i.e. without any direct link to the choice of clustering method. Below, we present a factorization method that performs clustering and marker gene identification in the same step.

Pseudotime, or pseudotemporal ordering of cells, describes a one-dimensional projection of single-cell data that is based on a measure of similarity between cells (e.g. a distance in gene expression space). In conjunction with pseudotime inference, by ordering cells, cell trajectories or lineages can be inferred that describe cell state transitions over (pseudo) time (15, 16). Two major classes of methods for the estimation of pseudotime and cell trajectories are: *i*) performing dimensionality reduction on the full data and then fitting principle curves to the cells in low-dimensional space; *ii*) constructing a graph for which cells are nodes and edges connect similar cells (can be done in high or low dimensional space), and then calculating the minimum spanning tree (MST) on this graph (17).

Of the class (*i*) methods: Monocle 2 (18) infers pseudotime using a principle curve generated by iteratively computing mappings between the a high-dimensional gene expression space and a low-dimensional counterpart. Pseudotime is then predicted by measuring the geodesic distance from each cell to a root cell. SLICER uses locally linear embedding for dimensionality reduction before constructing a minimum spanning tree (MST) on the low-dimensional space to infer trajectories: pseudotime is again computed by the geodesic distance (the root cell should be specified by the user) (19). DPT calculates transition probabilities between cells using a diffusion-like random walk, and pseudotime is then predicted by defining a distance across these random walks (with respect to a root cell) (20, 21). TSCAN (22) and Waterfall (23) employ similar strategies by first embedding data into low-dimensional space by dimensionality reduction, and then constructing a MST. Current methods in class (ii) include Wanderlust (24) and Wishbone (25): these construct a cell-cell graph and infer pseudotime by computing distances from each cell to a root cell, with the possibility of refining the pseudotime using waypoints. A recent method, scEpath, takes an alternative approach by inferring a single-cell energy landscape and using this to estimate transition probabilities between cell states, and thus cellular trajectories (26). In a similar vein, CellRouter uses flow/transportation networks to identify cell state transitions (27). For the whole family of methods for pseudotime inference (the mathematical foundations of which vary considerably, see (28) for review), experimental validation of the temporal ordering of cells remains difficult (15).

Lineage inference is challenging, and despite significant efforts to predict lineages between cells from RNA (or DNA (29)) sequencing, several challenges remain. Most of the pseudotime inference methods discussed above can also be used to predict lineage relationships, however inferring multiple branch points (i.e. more than two lineage branches) often remains beyond reach (30) (BioRxiv: https://www.biorxiv.org/content/early/2018/11/16/261768). Slingshot (31) can infer multiple branch points, but may require branch-specific (end point) information to do so. Monocle (18) is able to infer multiple branch points, however (similar to most trajectory inference methods) it identifies cell subpopulations separately from lineage relationships.

In addition to lineage inference, predicting cell-to-cell communication between single cells or cell subpopulations is a major unaddressed challenge. This is in part due to the challenge of integration: multiple inferred properties, including cell subpopulations, lineages, and marker genes, are all involved in cell-cell communication, thus inferring cell communication is a high-level task that will only be successful following coherent characterization of each of these constituent properties.

Here we present SoptSC in order to address these challenges and infer cell lineage and communication networks. SoptSC, similarity matrix-based optimization for single-cell data analysis, performs multiple inference tasks based on a cell-cell similarity matrix that we introduce. The cell-cell relationships learned via the similarity matrix define which cells are clustered together, and from these complex lineages with multiple branches can be reconstructed. In the same step (i.e. decomposition of the similarity matrix), SoptSC also predicts marker genes for each cell subpopulation and along pseudotime. Communication networks between single cells are inferred using a probability model based on cell-specific expression of relevant sets of ligands, receptors, and target genes. By combination of these networks with the inferred cell lineage, SoptSC is able to predict complex regulatory interactions governing developmental cell trajectories.

To validate the clustering performance of SoptSC, We apply it to nine published scRNA-seq datasets with verified cluster labels and compare the results with those of other current clustering methods; on the same data we assess the ability of SoptSC to infer the number of clusters from the data. To quantitatively measure the accuracy and robustness of pseudotemporal ordering, we apply SoptSC to data on early embryo development and embryonic stem cell differentiation: systems where the biological stages (and thus experimental time) are well-characterized. Here we compare SoptSC to DPT (21) and Monocle 2 (18). We go on to apply SoptSC to scRNA-seq data on skin regeneration (32) to investigate the cell states, marker genes specifying each state, lineage relationships, and the cell communication and crosstalk between cells (mediated by specific signaling pathways, e.g. Bmp, Tgf-*β* and Wnt). These results help to illustrate the coherence achieved by performing multiple tasks simultaneously. In addition, we apply SoptSC to two scRNA-seq datasets on hematopoiesis (33, 34) to study the cell states, lineage relationships, and cell-cell communication networks that emerge. Here we also provide a comparison of the results between the two datasets.

## 2 MATERIALS AND METHODS

The methods of SoptSC are centered on a cell-cell similarity matrix (*S*), which is learned from the original gene-cell data matrix using a low-rank representation model (38). Clustering is then performed by applying rank-*k* (for *k* clusters) non-negative matrix factorization (NMF) to *S* (Figure 1). A truncated consensus matrix is then constructed, and by analysis of the eigenvalue spectra of its associated graph Laplacian (see Methods), SoptSC predicts the number of clusters (*k*). Also given by the rank-*k* NMF is an ordered list of genes that mark for each cluster, determined by the weights associated with genes in each cluster. Thus we obtain clusters and their marker genes in the same optimization step.

**Figure 1.**
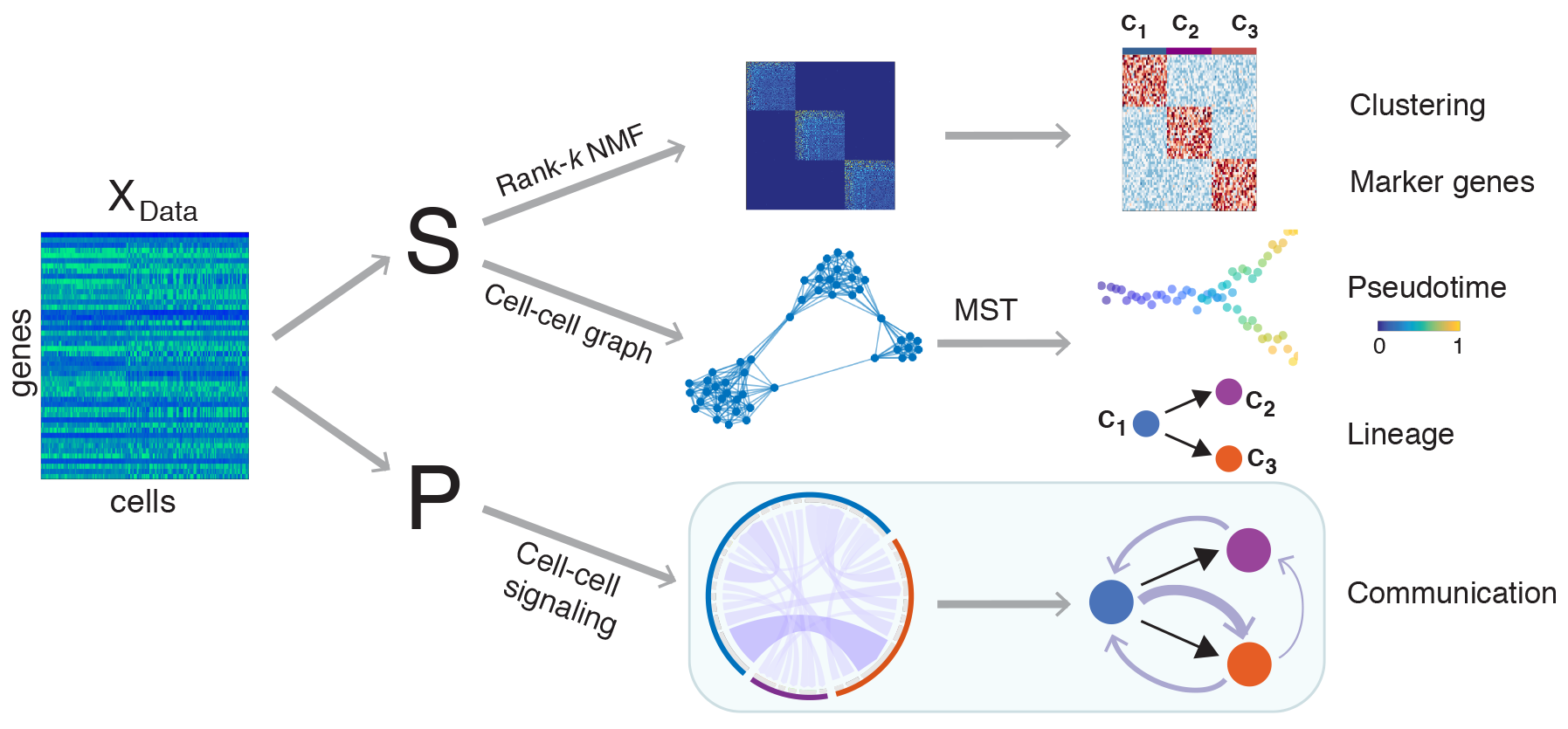
Overview of the SoptSC framework and outputs generated. SoptSC takes a gene expression matrix *X* as an input and learns a proper cell-to-cell similarity matrix *S*. Cell clustering is carried out by performing non-negative matrix factorization on *S*. Marker genes for each cluster are found via the product of the factorized latent matrix and *X*. A cell-cell graph (constructed from *S*) is used to infer pseudotime by calculating the shortest path distance between cells on this graph. The lineage relationships are constructed via a minimum spanning tree over the cluster-cluster graph derived from the cell-cell graph. Cell-cell communication is predicted by SoptSC via cell-cell signaling probabilities that are based on single-cell gene expression of specific genes within a pathway in sender-receiver cell pairs.

To infer pseudotime and cell lineage relationships, we construct a graph, the nodes of which are cells and the edges are given by the nonzero elements of *S* (See Methods). Using the cell-to-cell graph we define a distance as the shortest path length. A weighted cluster-to-cluster graph is constructed based on distances between constituent cell pairs, and the lineage hierarchy between subpopulations is inferred using the minimal spanning tree of the cluster-to-cluster graph. The longest path across the cluster-to-cluster graph identifies two end states. As these two clusters are far apart, it is often possible to identify which is the initial state via marker gene expression. Pseudotime is then inferred directly from the cell-to-cell graph by calculating distances between an initial cell and all other cells. The initial cell is chosen within the initial cluster such that pseudotime and the cluster-to-cluster (lineage) graph have the highest concordance.

In order to predict cell-cell communication networks (Figure 1), we identify signaling relationships based on single-cell gene expression for a pathway of interest. A signaling probability is defined based on weighted co-expression of signaling pathway activity in sender-receiver cell pairs (see details in Methods). As input, the user provides a ligand (or set of ligands) and cognate receptor (or set of receptors), for example: ligands from the Wnt family, and Frizzled receptors. For each pathway, a set of target genes is also specified: a candidate list of genes that are known to be differentially regulated downstream of a ligand-receptor interaction, along with their sign (upregulated or downregulated). SoptSC computes the signaling probability between sender cell (expressing ligand) and receiver cell (expressing receptor and exhibiting differential target gene activity). These single-cell signaling probabilities are combined to produce summaries and determine higher-level (e.g. cluster-to-cluster consensus) communication networks. Combining the consensus signaling networks with the lineage path allows SoptSC to infer feedback or feedforward interactions mediated by signaling factors.

### Cell-to-cell similarity matrix construction

The input to SoptSC is a single cell gene expression matrix *X* with *m* rows (associated with genes), and *n* columns (associated with cells). SoptSC computes the coefficient matrix *Z* from *X* by the following optimization model from Zhuang et al. (38):

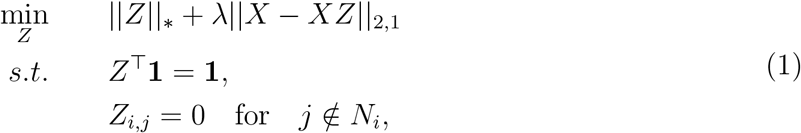

where || · ||_*_ is the nuclear norm; || · ||_2,1_ is the *l_2,1_* norm (the sum of the Euclidean norm of all columns); λ is a non-negative parameter, 1 = (1,…, 1)^T^ is a vector of length *n* and *N_i_* is the set of neighbors of cell *i*. To compute *N_i_*, cells are projected into low dimensional space using t-distributed stochastic neighbor embedding (t-SNE) (35), and a *K*-nearest neighbors (KNN) algorithm (36) is then applied to the low-dimensional data unlike in the previous study (35) in which KNN was directly applied without using dimension reduction. The linear constraint Z^T^1 = 1 guarantees translational invariance of the data (37).

The optimization model (1) is a representation method for the construction of graphs from nonlinear manifolds (38). Informally, it captures relationships between cells by representing each cell as a linear combination of all other cells. By restricting coefficients of non-neighboring cells to be zero, the model preserves the local structure of the linear representation. By imposing the low rank constraint, this model can capture the global structure of the original data input, and is more robust to noise and outliers. Problem (1) can be solved numerically by the alternating direction method of multipliers (38). Letting *Z*\* be the optimal solution of (1), then via symmetric weights we can define the similarity matrix as *S* = max { |Z* |, |Z*^T^|}. The elements *S_i,j_* (= *S_j,i_*) of *S* thus quantify the degree of similarity between cell *i* and cell *j*.

### Symmetric NMF for cell clustering

To cluster cells based on their similarity, we use symmetric non-negative matrix factorization (NMF) (39, 40) where the (non-negative) similarity matrix S is decomposed into the product of a non-negative low rank matrix 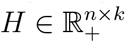 and its transpose *H*^T^ via the optimization problem:

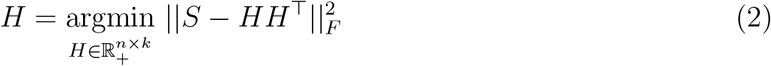

where || · ||_*F*_ is the Frobenius norm and *k* is the reduced rank of *S*. The solution of (2) satisfies the condition that the similarity matrix S can be represented approximately by *HH*^T^. Thus, the columns of *H* represent a basis for *S* in the (low rank) *k*-dimensional space, and the columns of *H*^T^ provide the coefficients for corresponding columns of *S*, in the *k*-dimensional space spanned by the columns of *H*. Since *H* ≥ 0, each column of *H*^T^ can be viewed as a distribution for which the *i^th^* column *S^i^* has the component in the corresponding column of *H*. We can use *H*^T^ to classify the N cells into *k* subpopulations by assigning the *i^th^* cell to the *j^th^* subpopulation when the largest element among all components of the *i^th^* column of *H*^T^ lies in the *j^th^* position. The clustering procedure based on the similarity matrix *S* is such that cells within a group have high similarity to each other and low similarity to cells from other groups. (Details can be found in supplementary materials).

### Marker gene identification

The non-negative low rank matrix derived from (2) not only produces cell cluster labels, but also marker genes for each cluster, by combining cluster labels with the data matrix in the following way. The element *H*_*i j*_· represents the weight by which cell *i* belongs to the *j^th^* cluster. Then the *j^th^* column of *H* defines a distribution over all cells corresponding to the *j^th^* cluster. It follows that we can define a weight for gene *v*, corresponding to the *j^th^* (1 ≤ *j* ≤ *k*) cluster, by:

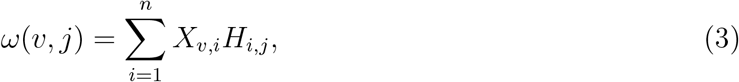

where *X_v,i_* represents the *v^th^* gene expression value in cell *i*. These weights for gene-cluster pairs measure the significance attributed to a given gene in each cluster, thus providing a means to determine how well gene *v* delineates cluster *j* from all other clusters. Marker genes are then defined as follows: gene *v* is a marker for cluster *j*, if *ω*(*v*,*j*) reaches its largest value in cluster *j*, i.e. *ω*(*v*,*j*) = max_1_≤u≤k {*ω*(*v*,*u*)}.

### Prediction of the number of clusters within a dataset

Prediction of the number of clusters is based on construction of a truncated consensus matrix and analysis of the spectrum of this matrix (41, 42). The number of clusters *k* is varied in a range {2, 3,…, *N*}, and symmetric NMF is then performed for each *k* to identify *k* clusters. A consensus matrix C is defined such that *C_i,j_* represents the number of times that cell *i* and cell *j* are classified as belonging to the same cluster. To improve robustness of this estimation to noise, we prune *C* by setting *C_i,j_* = 0 if *C_i,j_* ≤ *τ*(*N* — 1) where τ ∈ [0, 0.5] is the tolerance. The graph Laplacian 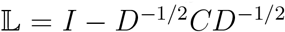 is computed, where *I* is the identity matrix, and *D* is a diagonal matrix of the row-sums of *C* (e.g., *D_i,i_* = Σ*_j_* C*_i,j_*).

We estimate the number of clusters within a range, by giving a lower bound as well as a upper bound. The lower bound is computed as the number of zero eigenvalues of 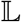, and the upper bound is computed as equal to the index at which the largest eigenvalue gap of 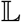 occurs. It has been shown that the number of eigenvalues of 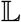 equal to 0 is equivalent to the number of diagonal blocks of 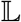 (41). By default we use the upper bound (the largest eigenvalue gap) as the estimate for the number of clusters. (Details can be found in supplementary materials).

### Inference of pseudotime and cell lineage

To infer the cell lineage and a temporal ordering of cells, we construct a cell-to-cell graph *G* = (*V, E, A*) where *V* is the vertex set of cells, and *E* is the edge set described by the adjacency matrix *A*. The adjacency matrix *A* is a binary matrix derived from the similarity matrix *S*: *A_i,j_* = 1 if *S_i,j_* > 0 and *A_i,j_* = 0 otherwise. We define the distance between cells on the graph *G* as the length of the shortest path between two cells on the graph, i.e. 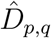 represents the length of the shortest path between cells *p* and *q* on the graph *G*.

Given a set of clusters *u* = {*u*_1_,*u*_2_, …,*u*_k_} (i.e. a partitioning of cells into *k* distinct groups), the distance between cell clusters *u_i_* and *u_j_* is computed by

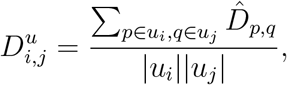

where |*u_i_*| represents the number of cells in cluster *u_i_*. 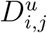 measures the average shortest path based distance between cluster *u_i_* and *u_j_*. A cluster-to-cluster graph *G^u^* = (*V^U^*,*E^U^*,*D^U^*) is constructed where each node in *V^u^* represents a cell cluster and *E^u^* is the edge set described by *D^u^*. The lineage tree, describing the cell state transition path, is inferred by computing the minimal spanning tree (MST) of graph *G^u^*. If the initial state is provided in advance, we construct MST by setting the root as the initial state. Otherwise, we provide prediction of the initial state (*u*_o_) by selecting the state that maximizes the path length over the MST.

Pseudotemporal ordering of cells is inferred by finding and sorting the shortest path lengths on the cell-to-cell graph *G* between each cell and the initial cell. An initial cell *c*_0_, if not provided in advance, is defined such that the temporal ordering of cells and lineage tree have the highest concordance. This is achieved as follows: for each cell *c_k_, k* ∈ *u*_0_ (i.e. cells clustered in the initial state), we compute 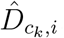 the shortest path length between *c_k_* and all other cells. By taking the mean shortest path length between *c_k_* and *c_l_*,*l* ∈ *u_i_* for each cluster *u_i_*, a mean distance from the initial cell to each cluster is defined. The Kendall rank correlation between the set of mean distances and the relative positions of states according to the lineage tree is then calculated. The initial cell is defined as that which produces the highest correlation by this test statistic.

### Pathway-mediated cell-cell signaling network inference

In order to study how paracrine signals are sent from and received by single cells, we implement a method to predict cell-cell signaling networks mediated by specific ligand-receptor interactions. Directed edges are drawn between two cells where a high probability of signaling is predicted by the expression of ligand in a “sender” cell, and the expression of its cognate receptor in a “receiver” cell along with appropriate expression of target genes of the pathway in the receiver cell. While such probabilities are not fully sufficient to define an interaction between a pair of cells, they represent necessary conditions for signaling, and can be indicative of spatial proximity of cells within a sample. Whereas previous works (43, 44) (BioRxiv: https://www.biorxiv.org/content/early/2017/09/27/191056) have considered signaling activity by summing over cells within a given cluster, we seek to account for the heterogeneity between cells within the same cluster.

For a given pathway, e.g. the Wnt signaling pathway, we define a set of ligands as the protein products of the Wnt gene family, and a set of receptors as the Frizzled (Fzd) proteins that bind Wnts. Also necessary as input to this signaling inference method are a set of target genes affected by Wnt along with their sign, i.e. upregulated in response to Wnt, or downregulated in response to Wnt. Currently, the sets of target genes used are small and defined manually; while automation of this step is possible, it would introduce another level of uncertainty to be accounted for.

The probability of signal passing between two cells is then computed as follows. Suppose we have a ligand-receptor pair, the expression of which is given by the distributions 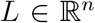 and 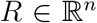 for ligand and receptor respectively, where *n* is the number of cells. Let 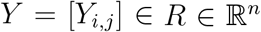 define the expression of *m*_1_ genes that are upregulated by Wnt, and 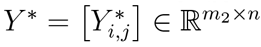 define the expression of *m*_2_ genes that are downregulated by Wnt. The probability that a signal is sent from cell *i* to cell *j* via this pathway is then given by:

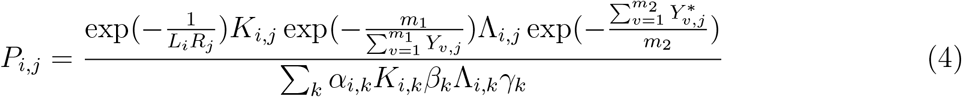

where:

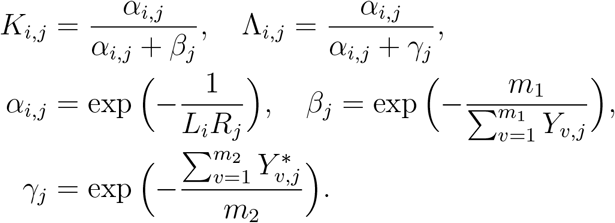

The first exponential term in equation (4) (*α_i,j_*) estimates the likelihood of an interaction between cell *i* and cell *j* given the expression level of ligand (*L_i_* in cell *i*) and the associated receptor (*R_j_* in cell *j*). If both are high, then the interaction probability is high; if either *L_i_* or *R_i_* is zero, the interaction rate is zero, i.e. there cannot be an interaction between cells *i* and *j* for this particular ligand-receptor pair. The second exponential term (*β_j_*) quantifies the expression of ‘activating’ target genes, i.e. those that are upregulated in cell *j* following a signaling cascade initiated by the ligand-receptor interaction. This term is weighted by the coefficient *K_i,j_*, which denotes that target genes can only increase the signaling probability if the likelihood of the ligand-receptor interaction is sufficiently large. Similarly, the third exponential term (*γ_j_*) quantifies the expression of ‘inhibitor’ target genes, i.e. those that downregulated the signaling pathway, thus decreases in inhibitor target gene expression correspond to increases in signaling probability. This term is weighted by the coefficient Λ_i,j,_ which acts similarly to consider the effect of inhibitor target genes only following ligand-receptor interaction. The signaling probability of a given edge is normalized by the sum of all possible signaling probabilities within the pathway.

The intuition underlying this formula is that if a ligand is highly expressed in cell *i*, the cognate receptor is highly expressed in cell *j*, and the corresponding target gene activity in cell *j* suggests that the signaling pathway may have been activated in this cell, then there is chance that that communication occurred between these two single cells, reflected by a higher signaling probability *P_i_,j*.

### Summaries of signaling networks and cluster—to-cluster signaling

Given a ligand-receptor pair for a specific signaling pathway, the signaling network inferred is given by the graph *G* = (*V, P*), where *v* is the set of all cells, and 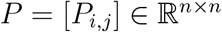 defines the probability of a signal being passed from cell *i* to cell *j* (Eqns. 4 and 5). For visualization of these networks we use the *circlize* package in R (45). A number of summary statistics derived from the probability matrix P can be defined, and are useful in various contexts.

### Consensus over signaling pathways

For a series of ligand-receptor pairs {*Lig_r_, Rec_r_*; *r* = 1, 2,…,*N*}, the corresponding signaling networks *G^r^* = (*V, P^r^*) are constructed based on each probability matrix *P^r^*, i.e. *P^r^* defines the probability matrix for ligand-receptor pair {Lig*_r_*, Rec*_r_*}. Then the overall probability of signaling summed over cells, *P^tot^* is given by:

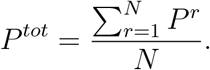

### Consensus over cells (cluster-to-cluster signaling)

It is also informative to consider the cluster-to-cluster signaling networks in order to predict where feedforward/feedback interactions may occur, and to compare with previous methods for cell-cell signaling study that have focussed on cluster-level signaling (43). Let *u* = {*u*_1_, *u*_2_,…, *u*_k_} give a clustering of cells by assigning each cell to one of *k* clusters. Then the probability of a signal passed between cluster *u_l_* and *u_m_*, mediated by a given ligand-receptor pair, is given by:

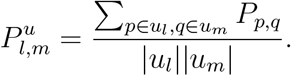

## 3 RESULTS

### SoptSC clusters cells in agreement with known identities

To assess the performance of SoptSC for clustering we compare it to four existing clustering methods: SC3 (10), SIMLR (9), Seurat (7), and tSNE (46) followed by k-means clustering (47) (tSNE + k-means) (Figure 2A), using nine published scRNA-seq datasets from a variety of biological systems (48–56). For each of these datasets, cell type cluster labels have been previously identified. Five of these are annotated as ‘gold-standard’ as the cell types have been verified by experiments; for the other four datasets labels were identified computationally (Table S1). Two of the gold-standard datasets (‘Deng’ (50) and ‘Pollen’ (49)) have two different possible clusterings: we test against both for each, giving a total of nine test cases. In order to measure the agreement between verified cluster labels and the cluster labels predicted by SoptSC, we use the Normalized Mutual Information (NMI) (57) as a test statistic. The value of NMI ranges from zero to one with higher value indicating higher accuracy in clustering, where one indicates perfect agreement between labels.

**Figure 2.**
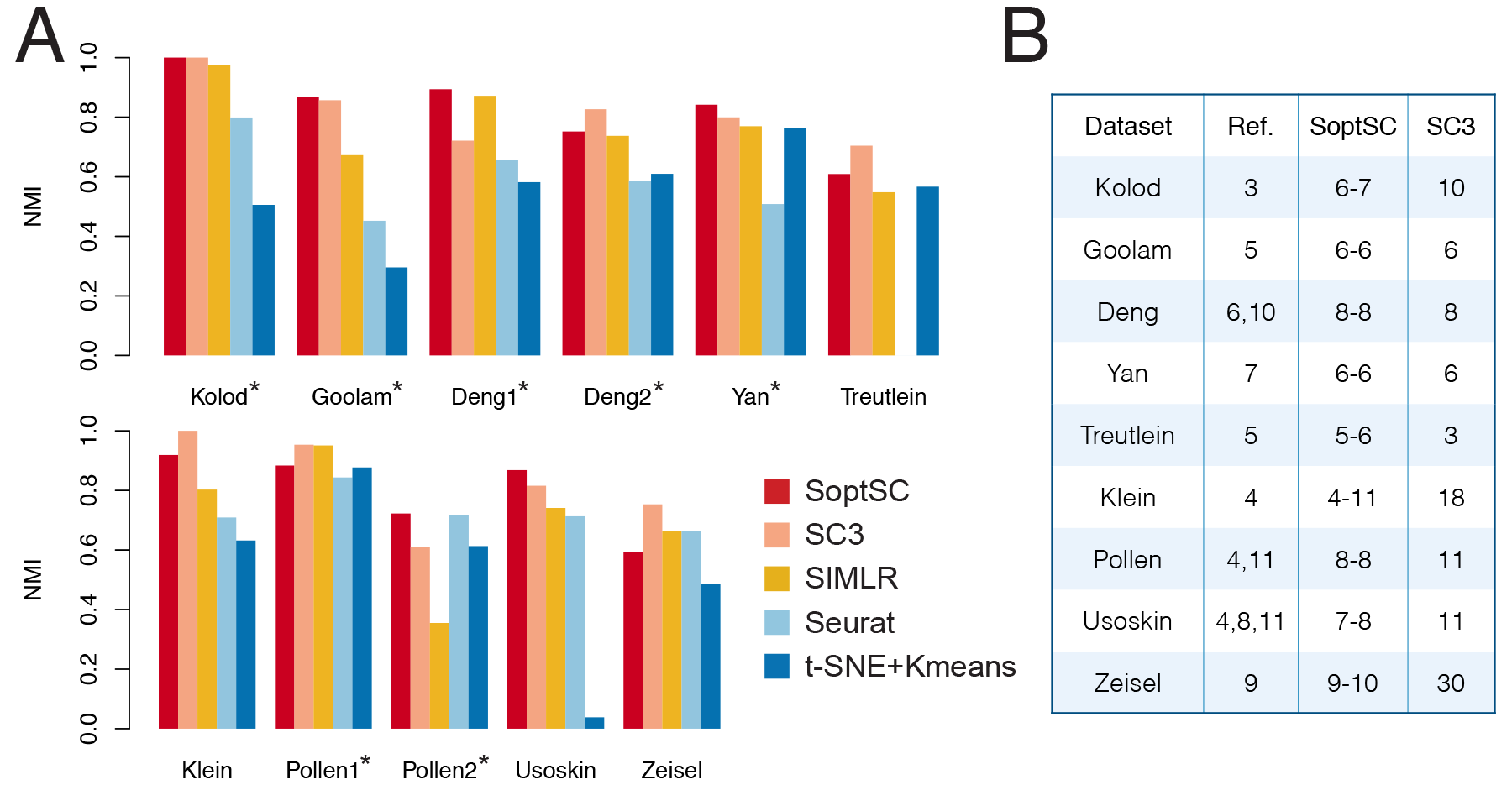
Benchmarking SoptSC against current methods for clustering. (**A**) Five clustering methods (SoptSC, SC3, SIMLR, Seurat, and t-SNE + Kmeans) are applied to a range of single-cell datasets where cell cluster labels are known or were previously validated. Normalized mutual information (NMI) is used as a measure of accuracy. Datasets marked by an asterisk are annotated ‘gold-standard’ for comparison purposes. (**B**) Prediction of the number of clusters by SoptSC or SC3, compared to a reference number of clusters (Ref.) from the original study; SoptSC predicts both lower and upper bounds.

For the three gold standard dataset with only one possible clustering, SoptSC has the highest NMI values across all methods compared. For the two datasets with two possible clusterings (Deng and Pollen), in each case, for one possible clustering SoptSC is best, and for the other possible clustering SC3 comes out on top. For the remaining four datasets, SoptSC shows comparable performance with SC3, and better performance than the other methods tested (Figure 2A). We also compare the number of clusters that SoptSC and SC3 predicts for each dataset. We also give the ‘true’ number of clusters, i.e. that of the original study, denoted as ‘Ref.’ (Figure 2B). In 6/9 cases, the number of clusters predicted by SoptSC was in agreement with the true number with a difference of no more than one. For SC3, this level of agreement was observed in 4/9 cases.

We thus find that SoptSC and SC3 exhibit superior performance to the other methods tested. Between these two methods, we found only slight differences between SoptSC and SC3. SoptSC outperforms SC3 for some datasets, and SC3 outperforms SoptSC for others; overall they are comparable for clustering (there seems to be a degree of complementarity between them on the datasets tested). In terms of in their ability to predict the number of clusters a priori from data, SoptSC outperforms SC3 on the datasets tested.

### SoptSC generates pseudotemporal orderings consistent with biological trajectories

Assessment of pseudotime inference is challenging as there are few examples of data for which the true ordering of cells is known or can be reliably estimated. One way to assess pseudotime is through its correlation with experimental time, e.g. using the Kendall rank correlation coefficient measured between pseudotime and the biological/experimental time labels obtained during data collection (21). Such a correlation can measure the ‘accuracy’ of pseudotime, with the caveats that this test can only be made at the (usually coarse) granularity of the experimental time points, and that cells are likely not synchronized in biological time. (Indeed, this is one of the reasons we infer pseudotime in the first place.) An estimate for the robustness of pseudotime inference can be made by subsampling of the data, i.e. we infer the pseudotemporal ordering of a portion (e.g. 90%) of the cells, and calculate the robustness as the correlation between the pseudotime of the subsample of cells and the pseudotime of the full data.

Using these two statistics as approximations for the pseudotime accuracy and robustness, we compared SoptSC to two current methods for pseudotemporal ordering: Diffusion Pseudotime (DPT) (21), which uses a diffusion map (a random walk over cells) to determine lowdimensional coordinates, and Monocle 2 (18), which uses reversed graph embedding to predict pseudotime. We test these methods using two datasets on development (i.e., early murine embryonic development (58) and murine embryonic stem cell differentiation (55)) and one dataset from bone-marrow-derived dendritic cells (59), which are chosen because of the clear temporal structure these systems exhibit, which should improve the reliability of the time correlation as a test statistic. We also used a dataset from murine interfollicular epidermis where cell stage was inferred from the original study (32). In addition, there are well-curated sets of temporal marker genes available for these systems.

For each dataset, we inferred the subpopulation structure and pseudotime via SoptSC and compared the pseudotemporal trajectories against the data and alternative methods. For all the datasets, the pseudotemporal ordering inferred by SoptSC was found to be highly consistent with known developmental stages (Figure 3A-B and Figure S2-S3). We also found that the accuracy of the pseudotime inference was higher for SoptSC than for DPT or Monocle2 for each of the datasets considered (Figure 3C-F). Under subsampling, SoptSC displays greater robustness than Monocle2, and is within 10% of DPT in terms of robustness (Figure 3C-F). These results indicate that, by the criteria used for assessment, SoptSC is more accurate - and comparable in robustness - as current state-of-the-art methods for pseudotime inference.

**Figure 3.**
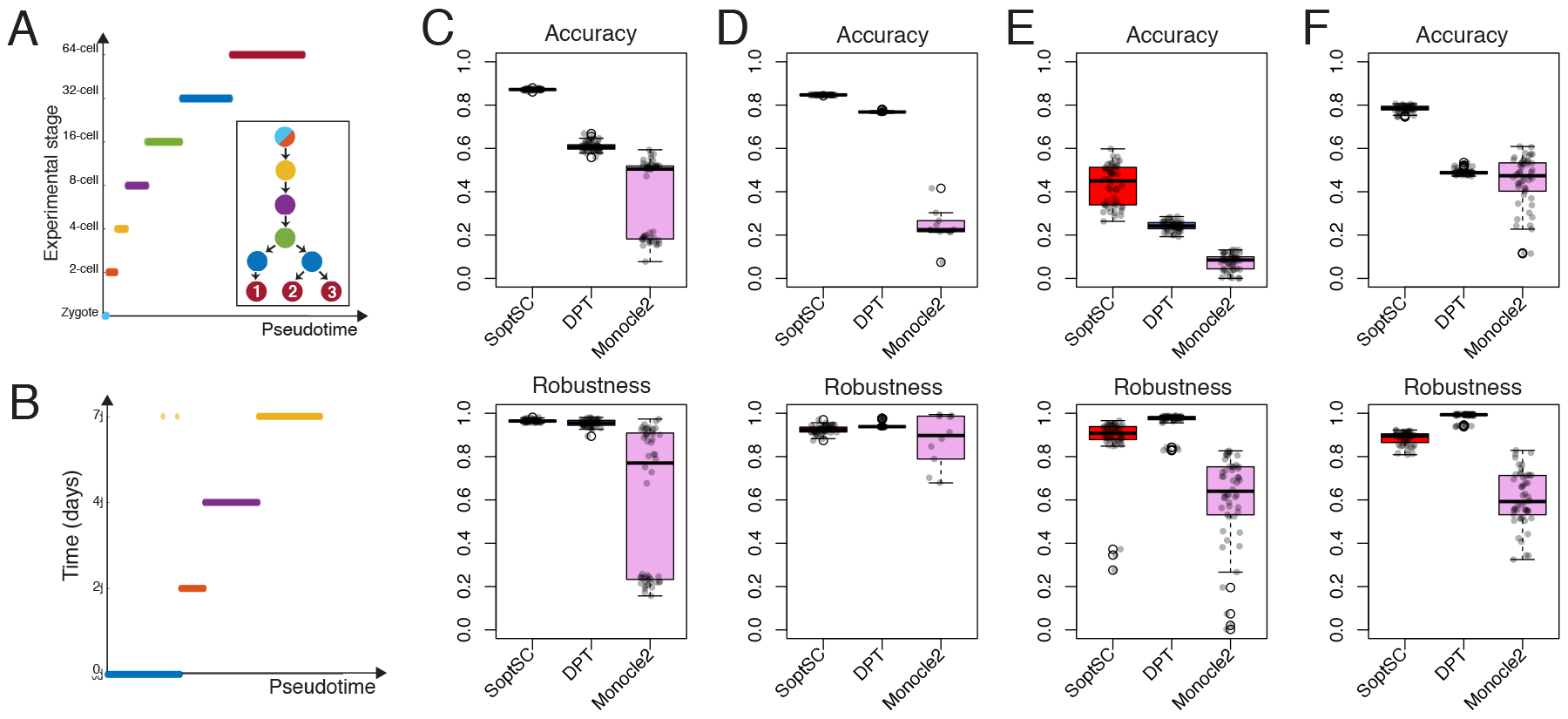
Assessment of SoptSC for pseudotime inference. (**A**) Pseudotemporal ordering of data from mouse early embryo development (58) is compared with the known biological stage. Inset shows the lineage inferred by SoptSC, colored by experimental stage of origin for each cluster. (**B**) Pseudotemporal ordering of embryonic stem cell data from (55) compared with experimental time. (**C**) Comparison of three methods for pseudotime inference with data from (58) using the Kendall rank correlation between pseudotime and experimental stage as a measure of accuracy, and by subsampling 90% of cells from the data 50 times (and comparison of subsets) to measure robustness. (**D**) Comparison as for (**C**) with embryonic stem cell data from (55). (**E**) Comparison as for (**C**) with bone-marrow-derived dendritic cells (59). (**F**) Comparison as for (**C**) with cells from the murine epidermis (32). Here the accuracy is measured by comparison with the pseudotime inferred in the original study.

### Inference of lineage hierarchies and marker gene dynamics recapitulate early embryonic trajectories

We analyzed two well-characterized embryonic datasets, for which, at least at early time points, the lineage relationships are well-characterized (48, 58). We use these data as further tests of clustering and pseudotemporal ordering in SoptSC, as well as to assess the inference of lineage hierarchies. We first look at the data from Guo et al. (58) (presented above in relation to pseudotime assessment), and find that SoptSC identifies nine cell subpopulations, which can be ascribed biological labels by comparison with known experimental stages (Figure 4A-B). The first of these subpopulations groups together zygotes and cells from the 2-cell stage. The second subpopulation almost exclusively contains cells from the 4-cell stage, and the third and fourth subpopulations contain 8-cell and 16-cell stage cells. Two subpopulations contain 32-cell stage cells (indicating a branch point here): analysis of marker genes identifies this branch point as giving rise to the trophectoderm (TE) and the inner cell mass (ICM) (Figure 4C, D). At the 64-cell stage, a second branch point is found emerging from the ICM, which forms (again, identified via marker genes) subpopulations representing the primitive endoderm (PE) and the epiblast (EPI). SoptSC was able to infer developmental stages from zygote to epiblast with high fidelity. The pseudotemporal cell ordering along this trajectory is also consistent with the developmental stages (Figure 4E). In addition, we have compared the clusters identified by SoptSC with alternative methods (SC3, Seurat, and SIMLR) and the inferred pseudotime (with Monocle2 and DPT). We found that SoptSC best recovers known biological information, both for clustering and pseudotime inference (Figure S2, S11).

**Figure 4.**
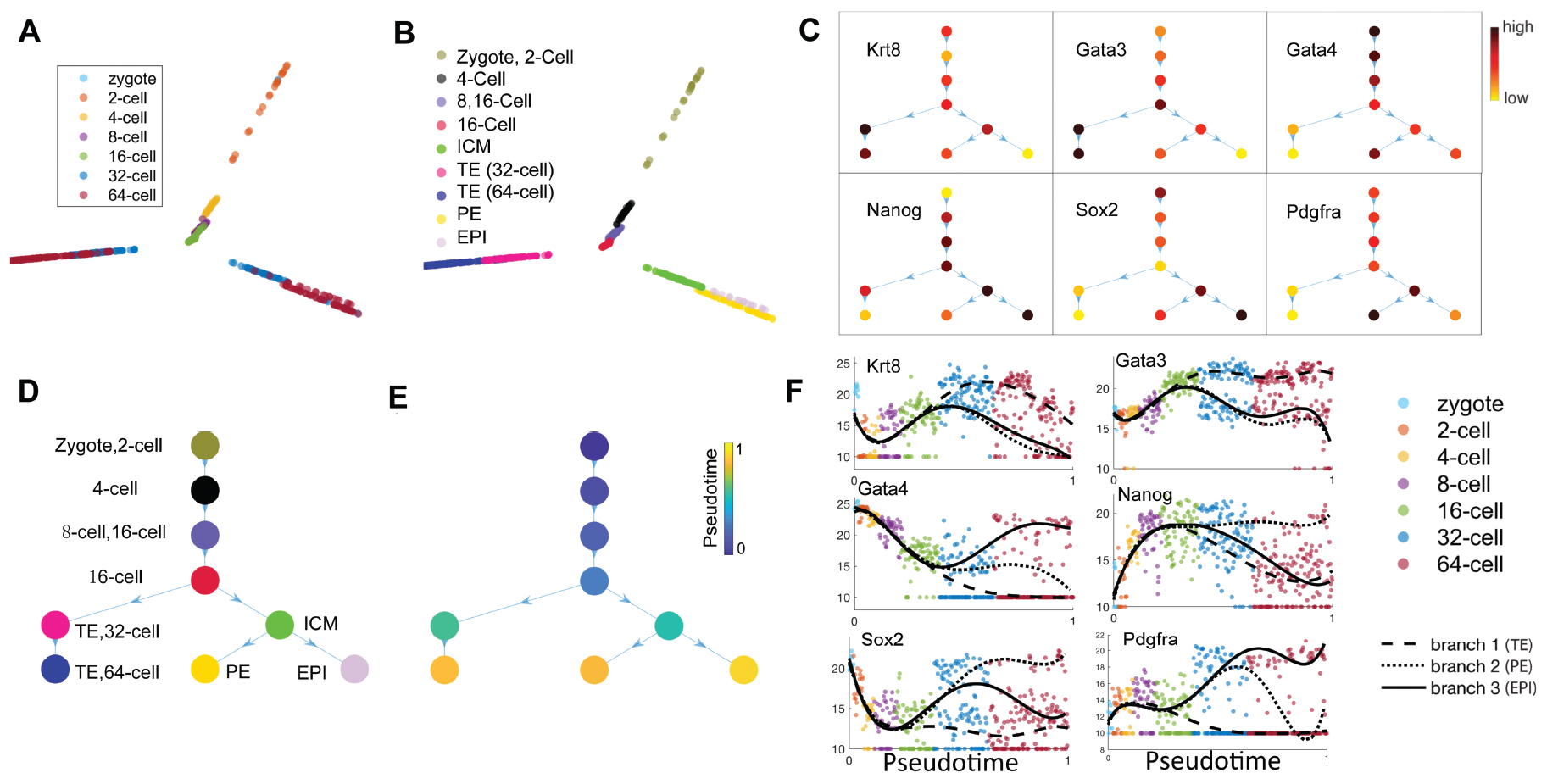
Inference of subpopulation, pseudotime and lineages for single cell data from mouse early embryos (58). (**A**) Visualization of data from mouse early embryo in two dimensions by SoptSC and colored by the experimental time stages from the original study. (**B**) Nine clusters were identified by SoptSC; labeled according to marker gene expression profiling. (**C**) Average expression of selected markers and plotted on the lineage tree inferred by SoptSC. (**D**) Lineage inferred by SoptSC with cluster labels, identified via known markers, from (C), and experimental stages, from (A). (**E**) Pseudotime projected onto the lineage tree. (**F**) Marker genes plotted along pseudotime; lines correspond to polynomial regression for each branch. TE: Trophectoderm; PE: Primitive Endoderm; EPI: Epiblast.

SoptSC can also resolve branch-specific marker gene dynamics along pseudotime. Six marker genes identified by SoptSC for the data from (58) are plotted along pseudotime in (Figure 4F), these show clearly distinguishable signatures for each lineage branch. By Gata4 alone, it is possible to identify all three lineages at a point late in pseudotime (around 32- to 64-cell stages): high expression marks cells of the epiblast; intermediate expression marks primitive endoderm, and low expression marks the trophectoderm.

We studied a human embryonic dataset with the similar goals of assessing inference of lineage and marker gene dynamics in SoptSC. For the data from Yan et al. (48), SoptSC identifies subpopulations corresponding to known development stages (Figure S8A, B, G), and a linear lineage trajectory from oocyte to late blastocyst, which is consistent with the inferred pseudotime (Figure S8C-F). We are encouraged that SoptSC was able to extract distinct developmental stages for even very few cells (only 88 cells in the dataset). To study gene dynamics along pseudotime, we plot six genes previously identified as embryonic markers (Figure S8H) and find good agreement between the dynamics predicted by these and previous studies (60).

### Temporal cell signaling activity regulates epidermal regeneration during telogen

The mammalian epidermis is a well-characterized adult stem cell system (61), yet significant questions regarding the constituents of epidermal cell subpopulations, the heterogeneity among them, and the interactions between them remain (62). Cells can (and in some cases must) transition between multiple states, such as in the case of hair follicle formation (63). In the interfollicular epidermis (IFE), cells are highly stratified: a stem cell population in the basal layer maintains the tissue through proliferation and production of differentiated cells; the keratinized cells form the outermost layer of the skin that is eventually shed (61).

We applied SoptSC to a murine epidermal dataset studying skin regeneration during the second telogen (32), in order to investigate the cell states present, their lineage relationships, and the signaling crosstalk between cells that regulates these relationships and their fates. Joost et al. (32) performed multi-level clustering in order to identify the various subpopulations of the epidermis, and found five subpopulations within the interfollicular epidermis (IFE) at the first level of clustering, one of which was basal (Figure 5A). At second-level clustering Joost et al. found heterogeneity — subpopulations — within the basal subpopulation. Here we focus our analysis on the IFE, as it likely represents a faithful trajectory in pseudotime, and we thus analyze 720 single cells. Within these data, we identify seven subpopulations, three of which are basal (clusters *C*_1_, *C*_4_, and *C*_5_, Figure 5B), thus, in contrast to the original analysis, we identify multiple basal subpopulations within the IFE in a single step. In addition, we compared the clusters identified by SoptSC with alternative methods (SC3, Seurat, and SIMLR) and the inferred pseudotime (with Monocle2 and DPT). We found that SoptSC best recovers known biological information, both for clustering and pseudotime inference (Figure 3F and Figure S4, S11).

**Figure 5.**
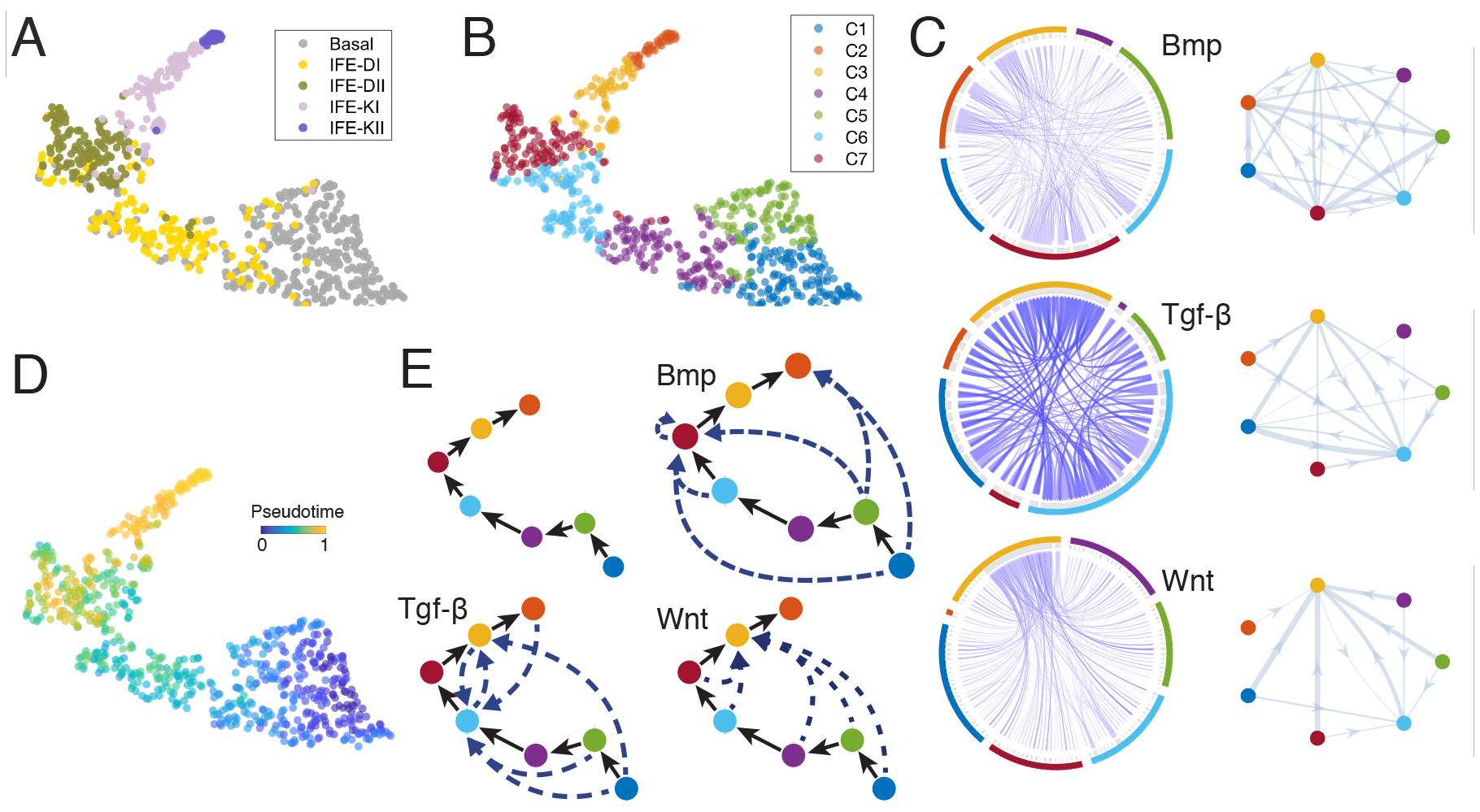
Inference of epidermal lineage and signaling networks. (**A**) Cells from (32) projected into low dimension by SoptSC and colored by the cluster labels from the original study. (**B**) Cells colored by cluster labels identified by SoptSC. (**C**) Single-cell communication networks predicted for three pathways. Left: samples from full networks where edge weights represent the probability of signaling between cells. Right: cluster-to-cluster signaling interactions where edge weights represent sums over within-cluster interactions. Colors correspond to cluster labels from part b. (**D**) Pseudotemporal ordering of cells. (**E**) SoptSC infers a linear lineage from basal to differentiated epidermal cells (top left). Summaries of the cluster-to-cluster signaling interactions with highest probability are given for the Bmp, Tgf-*β*, and Wnt pathways.

We study cell trajectories by inferring the pseudotemporal ordering of cells, the lineage relationships between cell subpopulations, and the means by which cells pass signals via cellcell communication (Figure 5C-E and Figure S7C and D). The pseudotime and lineage show a linear hierarchy between subpopulations, suggesting an initial (most stem-like) basal state gives rise to two restricted basal states before differentiating and eventually becoming keratinized (clusters *C*_2_, *C*_3_, *C*_6_, and *C*_7_). From the gene expression of six epidermal markers of the IFE (Figure S7B), we see broad agreement with known epidermal cell biology whereby basal cells appear earliest and must differentiate before eventually become keratinized (late in pseudotime). We also found that the marker predicted (unsupervised) by SoptSC overlapped considerably with known markers of epidermal differentiation (Figure S7F).

In order to study how cell-to-cell signaling regulates cell state transitions during epidermal differentiation, we construct single-cell communication networks defined by specific pathways. Here we study three pathways: Bmp, Tgf-*β*, and Wnt (Figure 5C and Figs. S12-S13). In Figure 5C we plot both the most probable signaling interactions between subpopulations (right), and between individual cells (left). While the subpopulation signaling plots provide useful and easily interpretable summaries, it is also important to study putative interactions between single cells, as these can be indicative of a physical proximity of two cells (providing at least a necessary condition), and will also become very important in downstream analyses, as we predict that the field will soon begin building up cell networks within tissues from the level of single cells. Crucially, in our predictions of cell-cell communication, we use expression of target genes of the pathway of focus as evidence that a signal to that cell is present; we currently use small curated lists of target genes for this purpose, but these can be readily grown by users with particular domain knowledge.

We observe that these pathways show distinct temporal patterns along the differentiation trajectory of keratinocytes. The highest probability of cells interacting through Bmp signaling is predicted to occur in mid-differentiated epidermal population *C*_7_, marked by Krt77 and Ptgs1. The Tgf-*β* pathway is activated in the subpopulations *C*_6_ and *C*_3_. These mark early differentiating cells, and late differentiated/early keratinized cells, respectively, thus indicating greater number of sites of influence due to Tgf-*β* signaling than Bmp. Experimental work has demonstrated that Tgf-*β* is crucial for the terminal differentiation of keratinocytes (64), thus supported our prediction. Wnt signaling is predicted to be active specifically in *C*_3_ (Figure 5C), the subpopulation undergoing keratinization; studies have shown that Wnt-Bmp signaling crosstalk regulates the development of mature keratinocytes, thus in agreement with these cellcell communication predictions in which neighboring differentiating epidermal populations are targeted by Bmp and Wnt (65).

The combination of these signaling networks predicts temporally constrained signaling pathway activation: Tgf-*β* being activated earliest during epidermal differentiation, followed by Bmp, and then Wnt, which is predicted to be important only subsequent to signals from the other two pathways. Given this prior knowledge that we have about the structure of the tissue and the locations of cell types within it, these analyses can prompt spatial pattern prediction from the signaling networks. Wnt signaling activity is predicted to be important only subsequent to signals from these other two pathways, thus suggesting that it acts farthest from the basal membrane.

### Hematopoietic lineage hierarchies coupled with single-cell communication reveal subpopulation-specific signaling interactions regulating differentiation

Hematopoiesis, the hierarchical formation of different blood cell types from a common multi-potent stem cell, involves complex cell state transitions and cell fate decision processes that are still incompletely understood (66–70). A multistep differentiation process involving successively restricted progenitor cell populations has been the dominant model to describe blood formation (71), although in some lineages restriction (to unipotency) may occur much earlier than previously thought (72). Significant heterogeneity within progenitor cell subpopulations is also leading to new perspectives on how hematopoietic cell differentiation proceeds (33, 73). The hematopoietic system thus provides an ideal for methods that seek to infer lineage hierarchical relationships; here we apply SoptSC to two such datasets to study the predicted lineage and signaling relationships (33, 34).

Olsson et al. (33) investigated myelopoiesis (hematopoietic differentiation restricted to myeloid lineages), and in particular found an intriguing cell state preceding the granulocyte/monocyte cell fate decision, containing cells of mixed lineage identity, which they named the ‘MultiLin’ cell state. We re-analyzed these data using SoptSC and found eight subpopulations (Figure 6A-B), which (via marker gene analysis) correspond well to known hematopoietic progenitor stages. We also inferred pathway-specific signaling networks (Figure 6C) and the hierarchical lineage relationships between subpopulations (Figure 6D). We have also compared the clusters identified by SoptSC with alternative methods (SC3, Seurat, and SIMLR) and the inferred pseudotime (with Monocle2 and DPT). We found that in each case SoptSC best recovers the biological information that is known about the system (Figure S10-S11).

**Figure 6.**
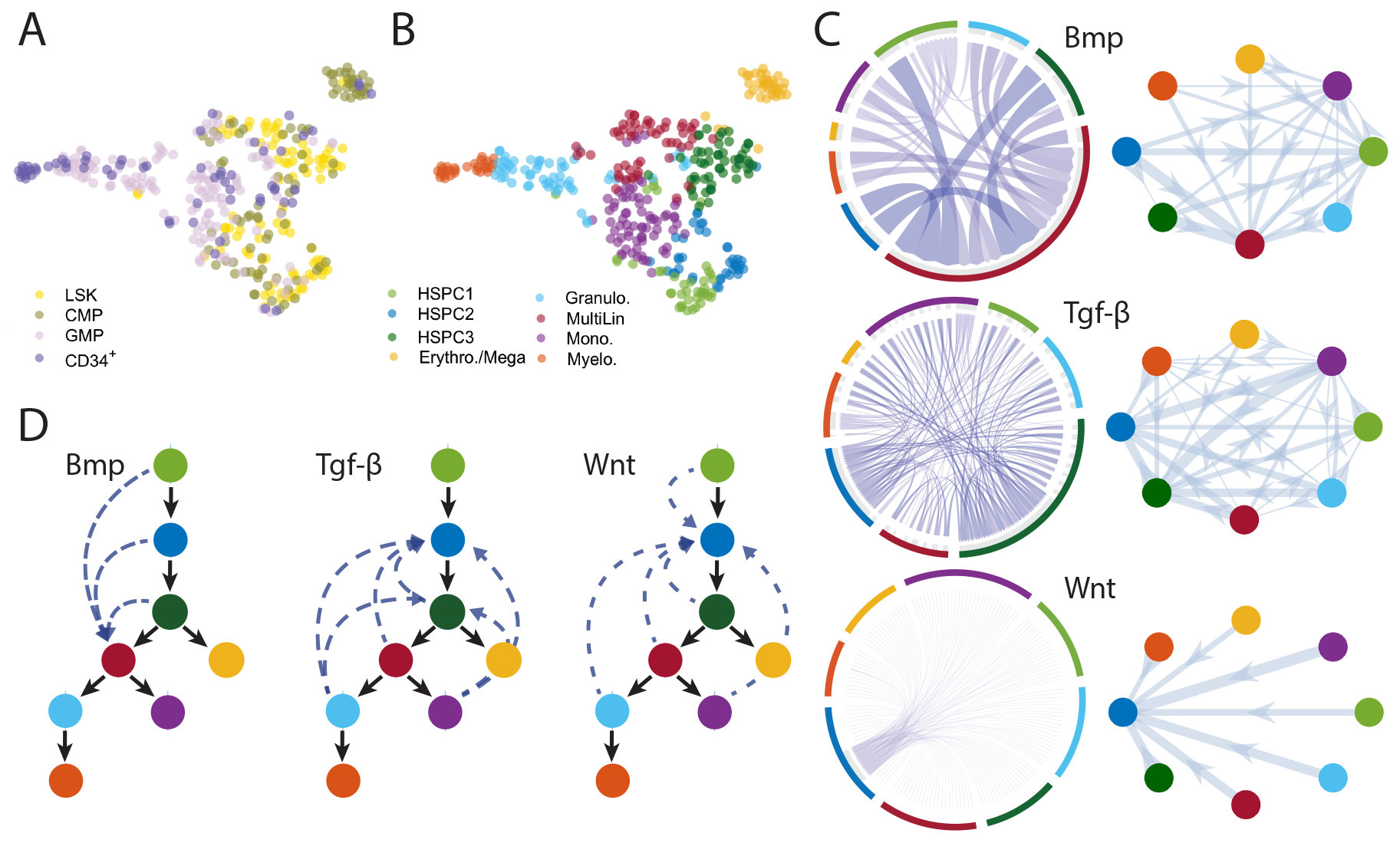
Inference of subpopulations, pseudotime, lineage paths and signaling networks during myelopoiesis. (**A**) Cells from (33) projected into low dimension by SoptSC and colored by the cluster labels from the original study. LSK: Lin^-^Sca1+c-Kit+; CMP: common myeloid progenitor; GMP: granulocyte monocyte progenitor; CD34+: LSK CD34+ cells. (**B**) Cells colored by cluster labels identified by SoptSC. (**C**) Single-cell communication networks predicted for three pathways. Left: samples from full networks where edge weights represent the probability of signaling between cells. Right: cluster-to-cluster signaling interactions where edge weights represent sums over within-cluster interactions. Colors correspond to cluster labels from part B. (**D**) Lineage inferred by SoptSC. Summaries of the cluster-to-cluster signaling interactions with highest probability are given for the Bmp, Tgf-*β*, and Wnt pathways.

The inferred lineage contained two branch points, the first of which gave rise to an erythrocyte/megakaryocyte progenitor subpopulation (marked yellow), and the second of which defined the granulocyte and monocyte lineages from a common progenitor (the ‘MultiLin’ state using the nomenclature of (33)). Notably, this lineage that contains multiple successive branch points and was inferred in a single step, contains greater information regarding lineage relationships than previous analyses of these data, which restricted inference of branch points to subsets of the data containing only one branch point (giving rise to granulocytes and monocytes) (18). Distinct gene signatures for the lineage branches were resolved along pseudotime via SoptSC (Figure S9E). The heterogeneity present during hematopoiesis is reflected in the marker gene expression heatmap (Figure S9D), in which certain subpopulations are difficult to distinguish, nonetheless, we find sufficient discriminative power for the genes that are identified by this method to be able identify the cell subtypes present. For example, Gata2 marks for erythrocytic progenitors, and Fos marks for hematopoietic stem/progenitor cells.

We next study the single-cell communication networks mediated by three pathways: Bmp, Tgf-*β*, and Wnt (Figure 6C-D), to investigate their effects on cell differentiation during myelopoiesis. The strongest effects due to both Wnt and Tgf-*β* are predicted to occur as feedback signals onto multipotent cell subpopulations. Experimental studies support the result that Wnt is most active during early hematopoiesis ((74) and references therein), however there are also controversies regarding the roles of Wnts during hematopoiesis (e.g. (75, 76)), highlighting the need for single-cell studies to disentangle these competing hypotheses. Tgf-*β* similarly regulates a large number of cellular processes, however it too is known to play a key role in the self renewal of hematopoietic stem cells (77), in agreement with the predictions of SoptSC. In addition we can compare our predictions with a ligand-receptor interaction database (44) that has been constructed (although with the caveat that Ramilowski et al. consider interactions in human cells). We find that this network predicts interactions between CD34+ and CD133+ stem/progenitor cells mediated by TGF-*β*, in agreement with SoptSC. These stem/progenitor cell populations (as well as myeloid progenitors) also express Fzd family ligands, thus making them candidate receivers of Wnt signaling, as our signaling network inference also predicts.

Bmp signaling is predicted to be most likely to be active in the MultiLin subpopulation (Figure 6D and Figure S14-S16). This mixed lineage state is particularly interesting as it defines a key transition immediately preceding the granulocytic/monocytic cell fate choice (33). From literature we find evidence that Bmp is crucial for maintaining the correct balance of myeloid cells, which lends support to our prediction of its activation in this myeloid progenitor subpopulation (78).

Within the multipotent/stem cell compartment, where we have predicted that three distinct subpopulations contribute to hematopoiesis (denoted HSPC1-3), we find that there is a very low probability of signaling (by any of these pathways) to HSPC1, the first hematopoietic cell state identified. This low-activity state is subsequently followed by HSPC2 and HSPC3, which are, by comparison, much more active in terms of individual cell-to-cell signaling mediated by all three ligands, and as such may represent activated/primed cell states prior to lineage restriction and commitment.

We also analyzed a hematopoietic single-cell gene expression dataset from Nesterowa et al. (34), which offers an opportunity to compare single cell hematopoiesis studies and assess the similarities and differences between them. For (34), SoptSC identifies six subpopulations (Figure 7A), which can be readily mapped to hematopoietic progenitor identities via marker gene expression (Figure S5D). We find for these data that a single subpopulation represents granulocyte and monocyte progenitors (GM), and that the dataset encompasses more of the hematopoietic tree, with distinct branches for lymphoid progenitors (L) and for erythrocyte and megakaryocyte progenitors (EM) (Figure 7B-C). The three branches inferred by SoptSC correspond closely to those identified in the original study: EM, GM, and L branches (see Figure 4A in (34) and Figure S5). The presence of the lymphoid progenitor cells in these data (and thus a wider spectrum of cell types) may lead to the granulocyte and monocyte subpopulations to be clustered together (the degree of granularity must always be chosen in light of the data at hand). We are also interested to discover what influence the lymphoid progenitor population has on the cell-cell signaling networks. In the case of this dataset, it was not possible to perform similar comparisons to the previous (Figure S4, S10-S11), since in this case no cluster labels were available with which to compare the predictions of SoptSC. However, given that we have already analyzed a hematopoietic dataset, we have the advantage that biological predictions made between the two can be directly compared.

**Figure 7.**
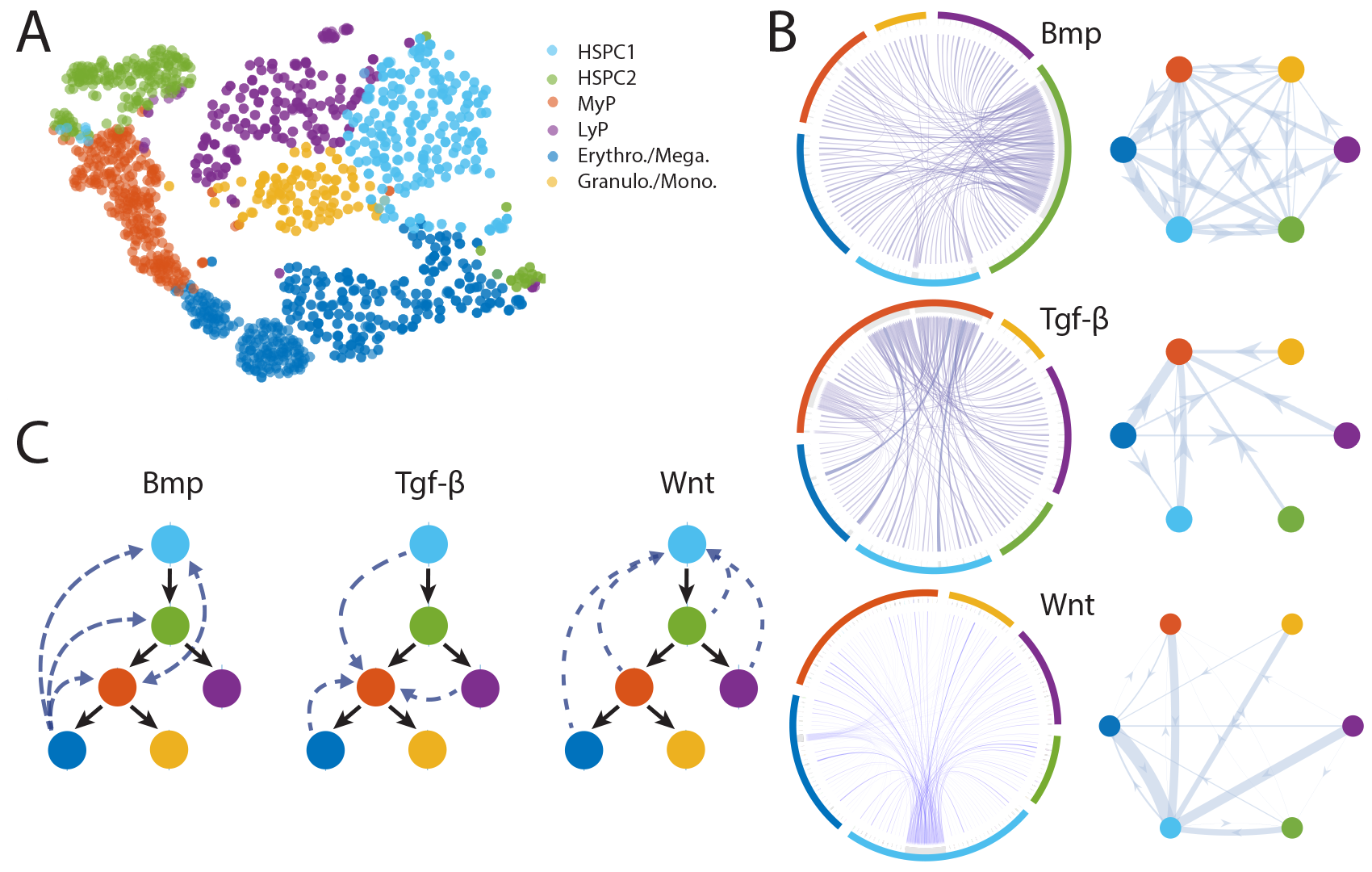
Inference of subpopulations, pseudotime, lineage paths and signaling networks for mouse hematopoietic stem cell differentiation. (**A**) Visualization and clustering of cells from HSPCs (34) by SoptSC. MyP: Myeloid Progenitor; LyP: Lymphoid Progenitor. (**B**) Single-cell signaling networks predicted for three pathways. Left: samples from full networks where edge weights represent the probability of signaling between cells. Right: cluster-to-cluster signaling interactions where edge weights represent sums over within-cluster interactions. Colors correspond to cluster labels from part A. (**C**) Lineage inferred by SoptSC. Summaries of the cluster-to-cluster signaling interactions with highest probability are given for the Bmp, Tgf-*β*, and Wnt pathways.

To infer signaling interactions and cell-cell communication, we employ the same methodology and use the same pathways as above, thus investigating Bmp, Tgf-*β*, and Wnt signaling (Figure 7B-C and Figure S17-S19). Overall, we see higher levels of ligand expression in each of the pathways than for the previous two datasets analyzed, and fewer ‘activated’ cells (cells that express receptor and regulate downstream target genes accordingly). During hematopoiesis as described in (34), Bmp signaling is predicted to provide feedback onto the multipotent progenitor populations from myeloid (GM) progenitors. There is also predicted to be activation within the myeloid progenitor population, in close agreement with the predictions made for (33) and (78). Wnt signaling is predicted to provide feedback onto the most naive stem cell population, which is again in agreement with the predictions from Olsson et al. and experimental work (74). Notably, the lymphoid progenitors are included in the set of subpopulations that activate via Wnt, providing signals that could improve the robustness of the myeloid-lymphoid branching cell fate decision (79). For Tgf-*β* signaling, there is some evidence of activation in stem cell populations, in agreement with Olsson et al., but the strongest predicted activation is in the myeloid progenitor population, suggesting that different feedback signals may be at play here, in part mediated by the lymphoid progenitors that are not present within the population of cells analyzed by Olsson et al.

## 4 DISCUSSION

We have presented SoptSC, an optimization-based method to infer subpopulations, cell lineage and communication networks from the high-dimensional single cell data in an unsupervised manner. At the heart of the method is a structured cell-to-cell similarity matrix, which is able to preserve intrinsic global and local structure of the data by incorporating low-rank constraints and allowing coefficients to be nonzero only in a local neighborhood of each data point (38). This local information is learned from the low-dimension representation of the gene expression data via t-distributed stochastic neighbor embedding, which is a key step for the construction of an appropriately structured cell-cell similarity matrix.

SoptSC provides prediction of the number of clusters present in a dataset by constructing the graph Laplacian of a truncated consensus matrix, and calculating the largest gap in its eigenvalue spectrum. SoptSC then concurrently clusters cells, infers pseudotime, marker genes, and a lineage tree from the similarity matrix. Clustering is achieved by performing NMF on the similarity matrix: the decomposed latent matrix is used to infer marker genes for each cluster in a way that is consistent with the clustering. We also implement a step to identify the initial cell in an unsupervised manner such that pseudotime has the highest concordance with the inferred cell lineage.

We have tested with favorable results the performance of SoptSC against current methods for single-cell data analysis against a variety of scRNA-seq datasets.

Importantly, the methods employed for inference of clusters, pseudotime, lineage, and marker genes derive from a single theoretical framework, making them directly and intuitively comparable. This is in contrast with most current scRNA-seq analysis pipelines (e.g. Seurat (7), SCANPY (80), and Monocle (18)), which combine multiple methods to analyze data. Seurat, for example, uses distinct methods to cluster cells and to find marker genes by identifying the most differentially expressed genes in cell subpopulations. SoptSC selects marker genes by identifying most likely expressed genes in each cell subpopulation, and consequently (in contrast with Seurat), SoptSC infers unique marker genes for each cluster. To study large, complex, and heterogeneous lineage-derived systems, it may well be worthwhile to perform multiple analyses by complementary means and seek the consensus between them. This can equally be helpful for studying and comparing pseudotime inference, as a recent large comparative study shows (BioRxiv: https://www.biorxiv.org/content/early/2018/03/05/276907).

The combination of cell lineage and cell communication predictions in SoptSC greatly facilitates biological discovery by identifying how particular cell state transitions can be regulated, by specific pathways, at the level of cell-to-cell communication. These are particularly relevant for regulatory interactions that occur during development or differentiation. For example, during hematopoiesis, analysis via SoptSC predicted that myeloid progenitor cell populations were targeted specifically by Bmp signaling for two datasets, whereas target populations of Wnt or Tgf-*β* signaling were more varied. During epidermal regeneration, we found temporally constricted signaling at play during keratinocyte differentiation, with Tgf-*β* signaling earliest to basal cells within the epidermis, before subsequent cell-cell communication by Bmp and then Wnt.

Recent studies have made significant progress in predicting gene regulatory networks from single-cell data (81, 82). SoptSC could also be combined with optimization approaches for gene regulatory network construction (83). Remaining significant challenges for scRNA-seq data analysis include confounding biological effects (e.g. due to cell cycle stages) and effects due to dropout (84, 85). Complementary methods developed to address these can be directly applied to input data as a preprocessing step before using SoptSC. For example, a recent method for imputation to handle gene dropout effects seems to be able to enhance scRNA-seq data quality (85), and several methods to regress out the effects of the cell cycle (86, 87) could also ameliorate data preprocessing. In addition, a recent method provides a statistical model that predicts which transcripts are likely to be affected by dropout noise, and which are not (85). This can be very helpful in identifying, e.g., ligands or receptors that are likely affected by dropout, and then to perform imputation on these genes specifically (rather than on the dataset as a whole); this helps to remove effects due to technical variation while preserving biological variation across cells.

Validation of new cell subpopulations predicted by scRNA-seq analyses is achieved first by successfully demonstrating that cells of this new subpopulation can be marked in vivo, e.g. via in situ hybridization with predicted marker genes. To ascribe function however, experiments such as clonogenic/differentiation assays, other culture following cell sorting by flow/mass cytometry, or genetic perturbations or target genes within the new subpopulation are required. New predictions provided by SoptSC of cell-to-cell communication patterns also must be rigorously tested in vivo. To do so, cells isolated from putative subpopulations (by cell sorting) could be plated and their responses to extraneous ligand tested. Subsequently, genetic perturbations to members (e.g. receptor) of a given pathway can provide further validation. However, we should also make full use of the dense information contained within a scRNA-seq dataset: we have predictions of communication not only between subpopulations but also between single cells. In analyses further downstream, this information will allow us to probe new cell states in higher resolution, e.g. by analyzing individual transcriptional states of different ‘signaling’ cells (either sending or receiving) within different cell subpopulations. We must be cautious of confounding effects here due to technical noise (see discussion of dropout above), but simultaneously signaling pathway analyses may provide additional means to handle dropout: i.e. it may be possible to integrate information about genes subject to dropout with information about pathways co-expression (from curated sets of members of a signaling pathway), as a way to improve the predictions of effects due to dropout, and thus the imputation process.

As datasets of 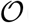(10^5^) cells become widespread (69, 88, 89), computational efficiency becomes a major challenge for many current single-cell data analysis methods, and new approaches are needed (90). For SoptSC, the computational cost associated with NMF contributes the most to the overall computational cost, and as such improvements and better algorithms for NMF will significantly improve the efficiency overall (91).

Lineage inference for complex heterogeneous data remains to be a challenging task. Compounding this is the lack of data for which the ground truth is known; since in many cases even the true cell states are not known, correct annotation of the relationships between is infeasible. SoptSC has nonetheless managed to infer complex lineage hierarchies that are correct to the best of current knowledge (such as for hematopoiesis). If we compare the lineage predicted by SoptSC for Olsson et al. data (33) with previous analysis of these data in (18), we find that unlike the previous analysis, in SoptSC are we able to resolve multiple branch points simultaneous. Nonetheless important improvements in lineage inference are needed. These include the ability to automatically detect multiple disconnected lineages within data, or to infer bidirectional arrows for some cell state transitions. Both of these features require an alternative fundamental approach to be taken, since the minimal spanning tree-based methods used for tree construction in SoptSC as well as in most current methods for lineage inference (18, 31, 92, 93) do not permit such features.

Single-cell data analysis comes with a particular set of promises and pitfalls. The key strength of scRNA-seq data lies in its ability to measure many thousands of signals simultaneously and provide a global quantification of the transcriptional state of a cell; this global cell state information however comes at the expense of accuracy in measurements of individual transcripts, due to noise and technical effects. With these challenges in mind, we have developed methods to perform multiple single-cell data analysis tasks coherently and unsupervised, and while much remains to be done, these have enabled new predictions of important cellular relationships, given by their positions within lineage hierarchies and communication networks.

### Software availability

SoptSC is available on GitHub as a MATLAB (Naticks, MA) package at: https://github.com/WangShuxiong/SoptSC. SoptSC is available as an R package at: https://mkarikom.github.io/RSoptSC.

### Availability of data and materials

All the datasets used in this paper are collected from the published accession numbers provided in the original studies. The datasets used for evaluating the performance of clustering (Figure 2) are downloaded from https://hemberg-lab.github.io/scRNA.seq.datasets/. Other datasets are available from: mouse early embryo (58) (Table S4): www.sciencedirect.com/science/article/pii/S15345 human early embryo (48): GSE36552; adult skin maintenance (32): GSE67602; myelopoiesis (33): GSE70245; data from Nesterowa (34): GSE81682; data from bone-marrow-derived dedritic cells (59): GSE48968.

### Author contributions

S.W., A.L.M. and Q.N. conceived the project. S.W. developed and implemented algorithms and software in MATLAB. M.K. developed software in R. S.W. and A.L.M. performed data analysis. S.W., A.L.M. and Q.N. wrote the paper; all authors read and approved the final manuscript. S.W. wrote the supplement. A.L.M. and Q.N. supervised the research.

## Supporting information

SoptSC_Supplement

## 5 ACKNOWLEDGEMENTS

We thank H. Singh for helpful discussions on hematopoiesis and on the data described in Olsson et al. (33).

