## Supplementary material for "Cell Lineage and Communication Network Inference via Optimization for Single-cell Transcriptomics": SoptSC_Supplement

### 1 Supplementary Figures

#### 1.1 Supplementary Figure S1

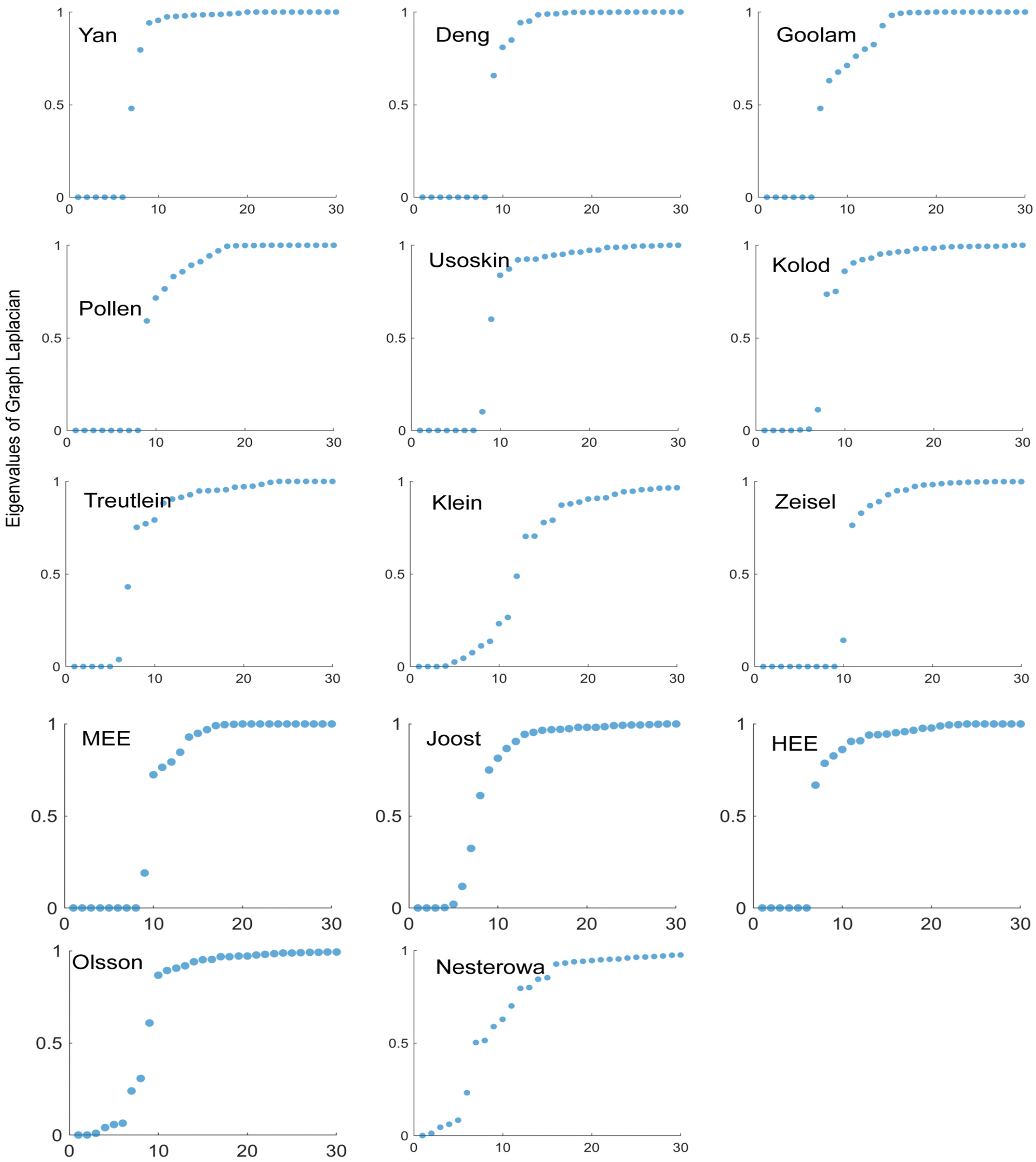

**Supplementary Figure S1. Spectra of the graph Laplacian of the truncated consensus matrix predict the number of clusters.** The first 30 sorted eigenvalues of the graph Laplacian of the constructed consensus matrix for each data set is plotted.

#### 1.2 Supplementary Figure S2

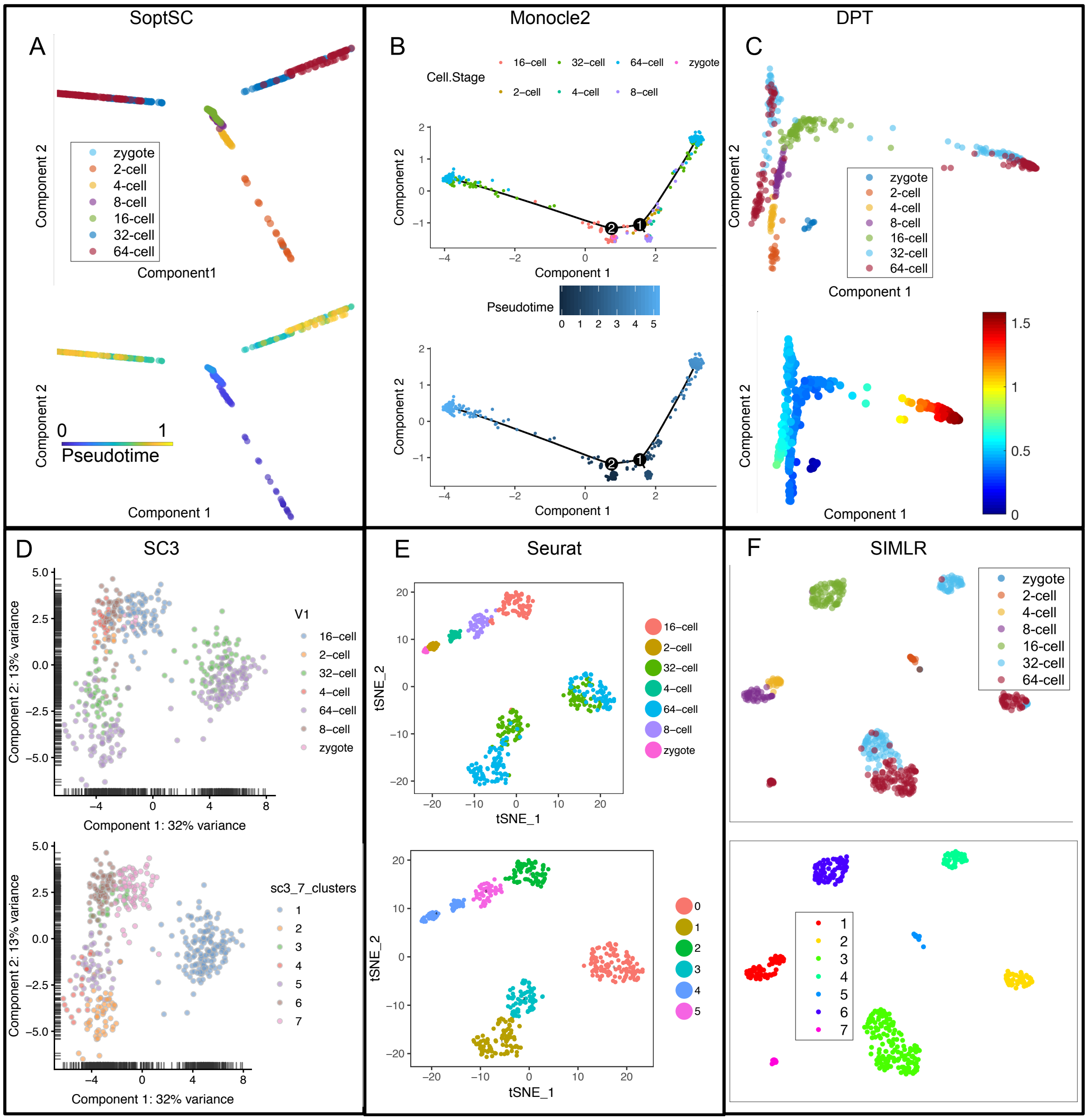

**Supplementary Figure S2. Pseudotime inference by SoptSC, Monocle2 and DPT; as well as clusters identified by SC3, Seurat, and SIMLR for mouse early embryonic data [4].** (A,B,C) Visualization of 2-dimensional trajectory of cells by (SoptSC, Monocle2, DPT) with true experimental time labels and pseudotime inferred by (SoptSC, Monocle2, DPT). (D,E,F) Visualization of low-dimensional projection of cells by (SC3,Seurat,SIMLR) with cell-stage labels and cluster labels identified by (SC3,Seurat,SIMLR).

#### 1.3 Supplementary Figure S3

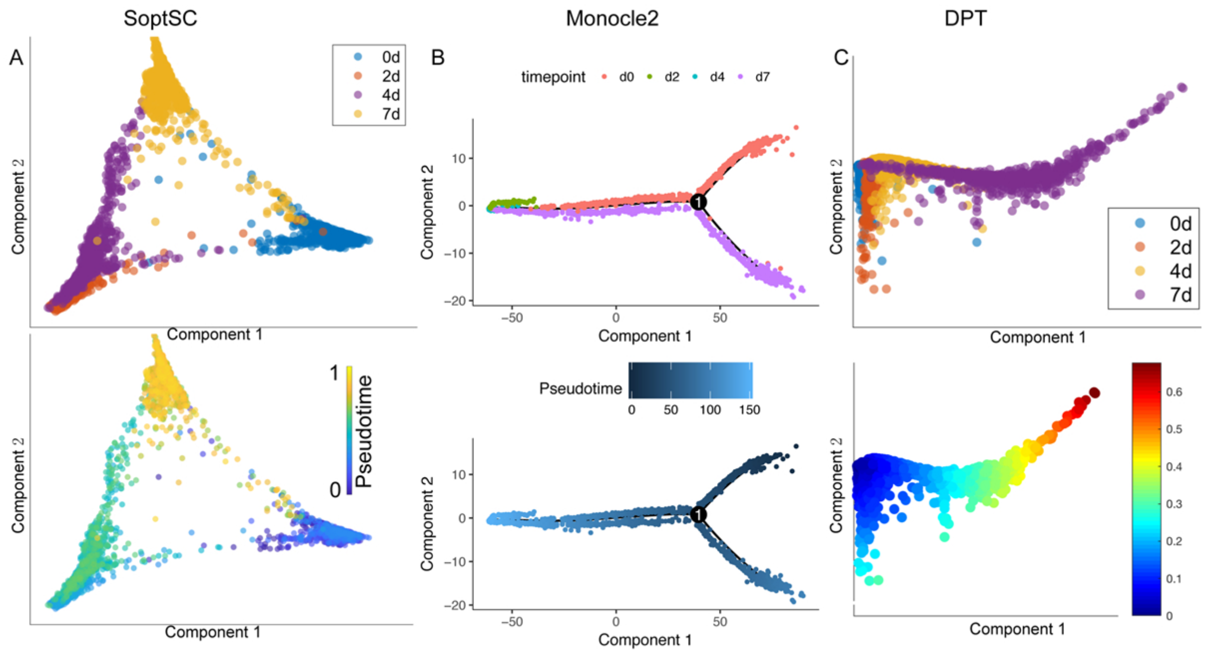

**Supplementary Figure S3. Pseudotime inference by SoptSC, Monocle2 and DPT for ESC data.** [7] (A) Visualization of 2-dimensional projection of cells by SoptSC with true experimental time labels and pseudotime inferred by SoptSC. (B) Visualization of low-dimensional trajectory identified by Monocle2 with true time labels and pseudotime inferred by Monocle2. (C) Visualization of low-dimensional projection of cells by DPT with true time labels and pseudotime inferred by DPT.

#### 1.4 Supplementary Figure S4

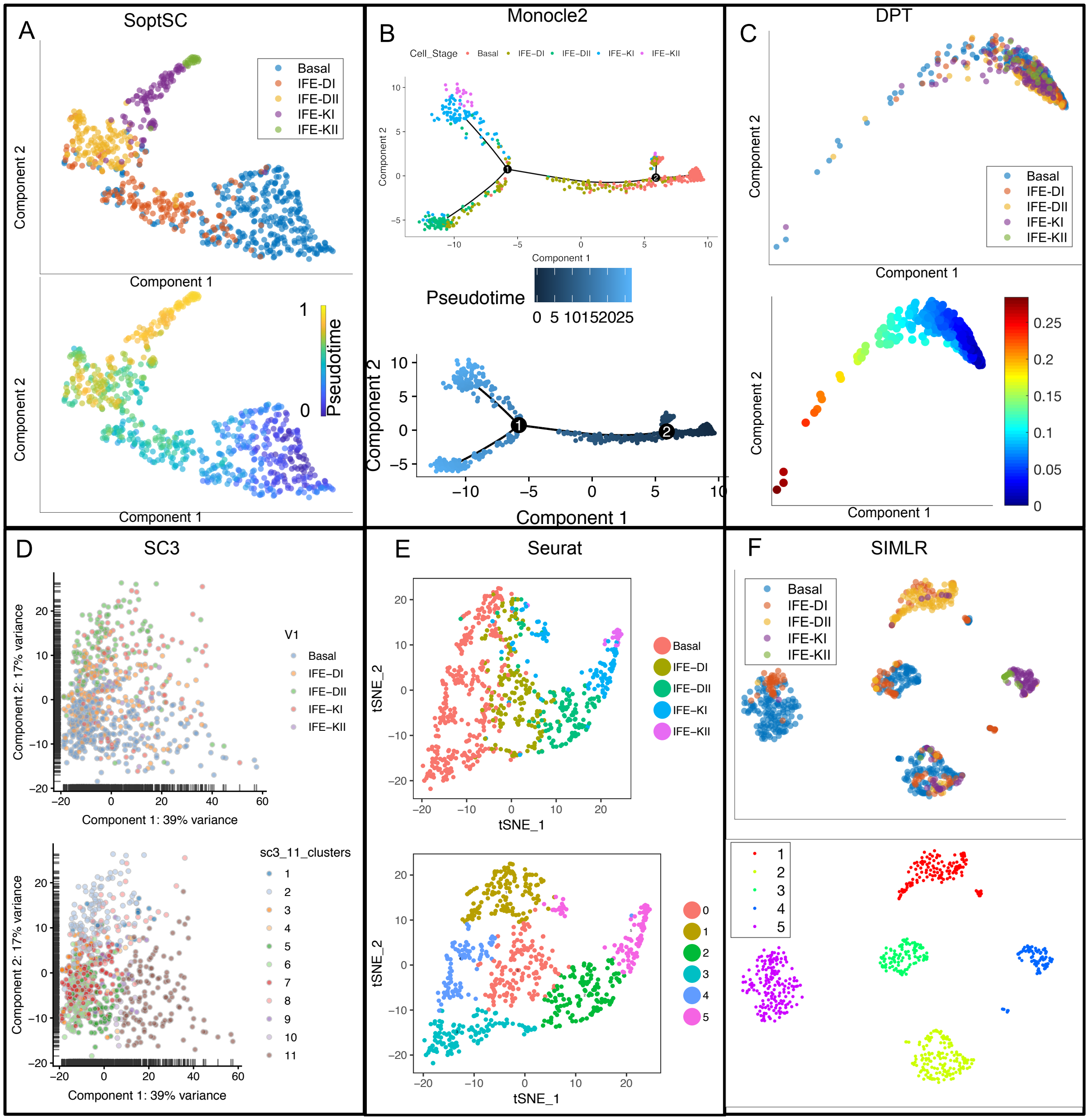

**Supplementary Figure S4. Pseudotime inference by SoptSC, Monocle2 and DPT; and clusters identified by SC3, Seurat, and SIMLR for IFE data [5].** (A, B, C) Visualization of two-dimensional projection of cells by (SoptSC, Monocle2, DPT) with true experimental time labels and pseudotime inferred by (SoptSC, Monocle2, DPT). (D, E, F) Visualization of low-dimensional projection of cells by (SC3,Seurat,SIMLR) with cell-stage labels and cluster labels identified by (SC3,Seurat,SIMLR).

##### 1.5 Supplementary Figure S5

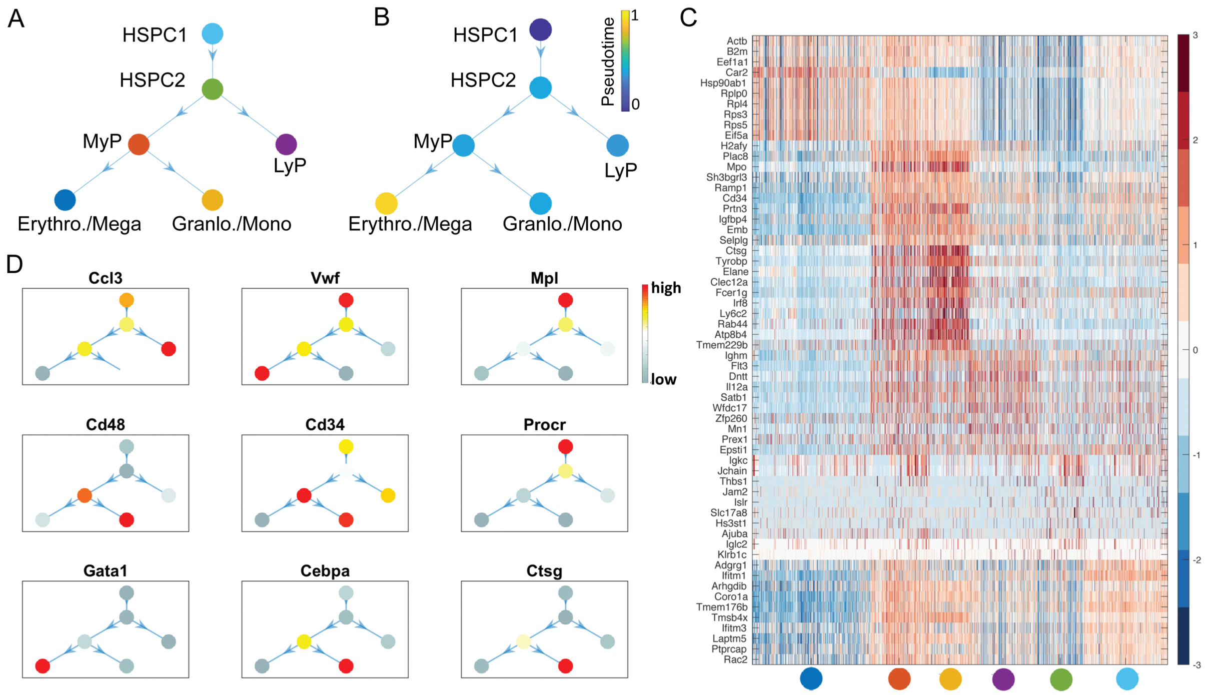

**Supplementary Figure S5. Analysis of pseudotime, and lineage paths for mouse hematopoietic stem cell differentiation [14].** (A) Lineage inferred by SoptSC. (B) Lineage hierarchy constructed by SoptSC. Colors correspond to the mean pseudotime value for the subpopulation. (C) Clustered gene-cell heatmap of genes from top 10 markers for each cluster identified by SoptSC; (D) Gene expression of selected markers. Colors represent the mean expression for each gene within each cluster.

#### 1.6 Supplementary Figure S6

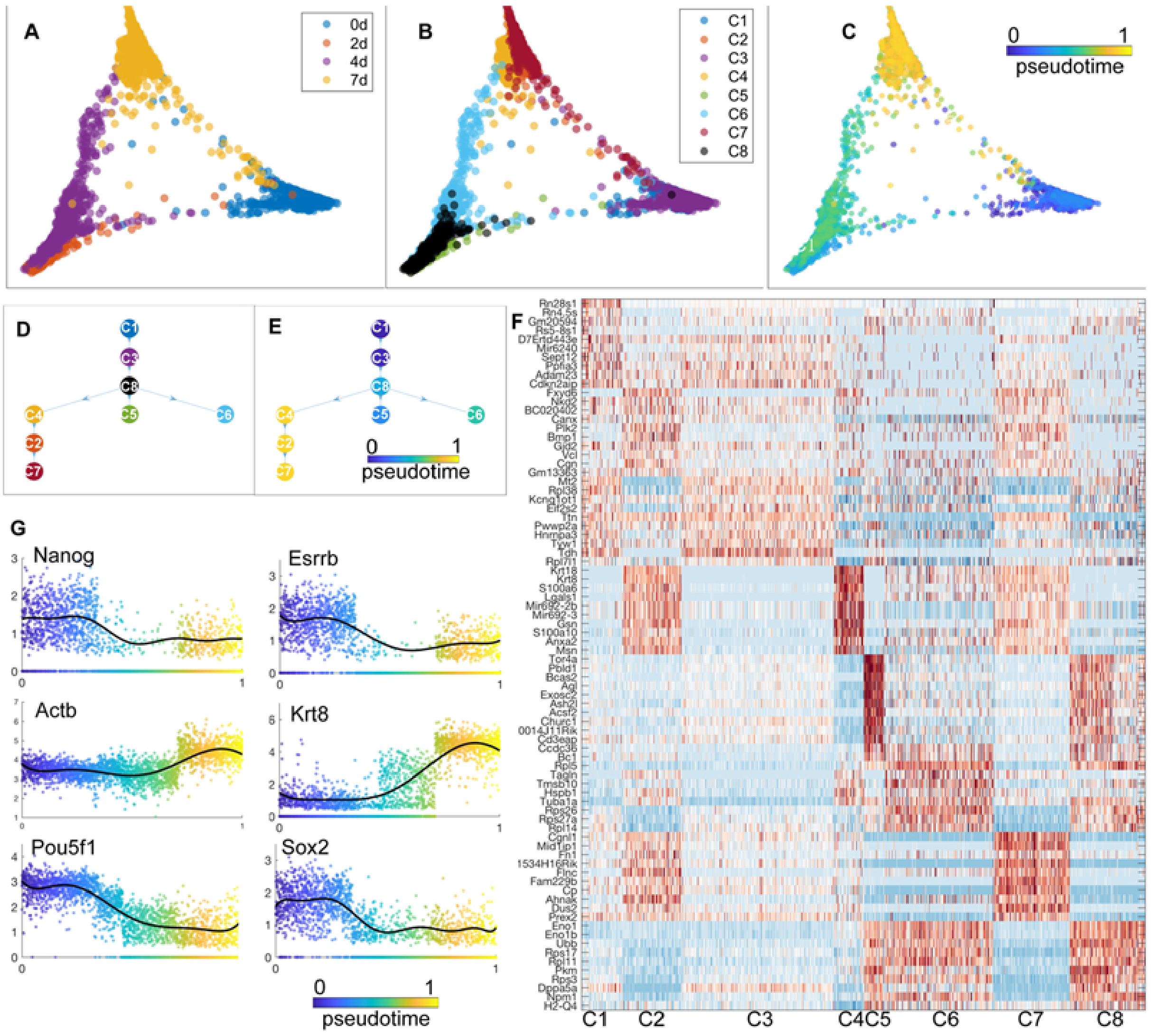

**Supplementary Figure S6. Analysis of the developmental trajectories of mouse ESCs [7].** (A) Visualization of cells labeled by true experimental time. (B) Cell subpopulations inferred by SoptSC. (C) Pseudotime for all cells. (D) Lineage inferred by SoptSC. (E) Pseudotime for cell states where the pseudotime of each state is calculated by the average of the temporal ordering of cells within the state. (F) Clustered gene-cell heatmap of genes from top 10 markers for each cluster identified by SoptSC. (G) Expression of selected marker genes along pseudotime.

#### 1.7 Supplementary Figure S7

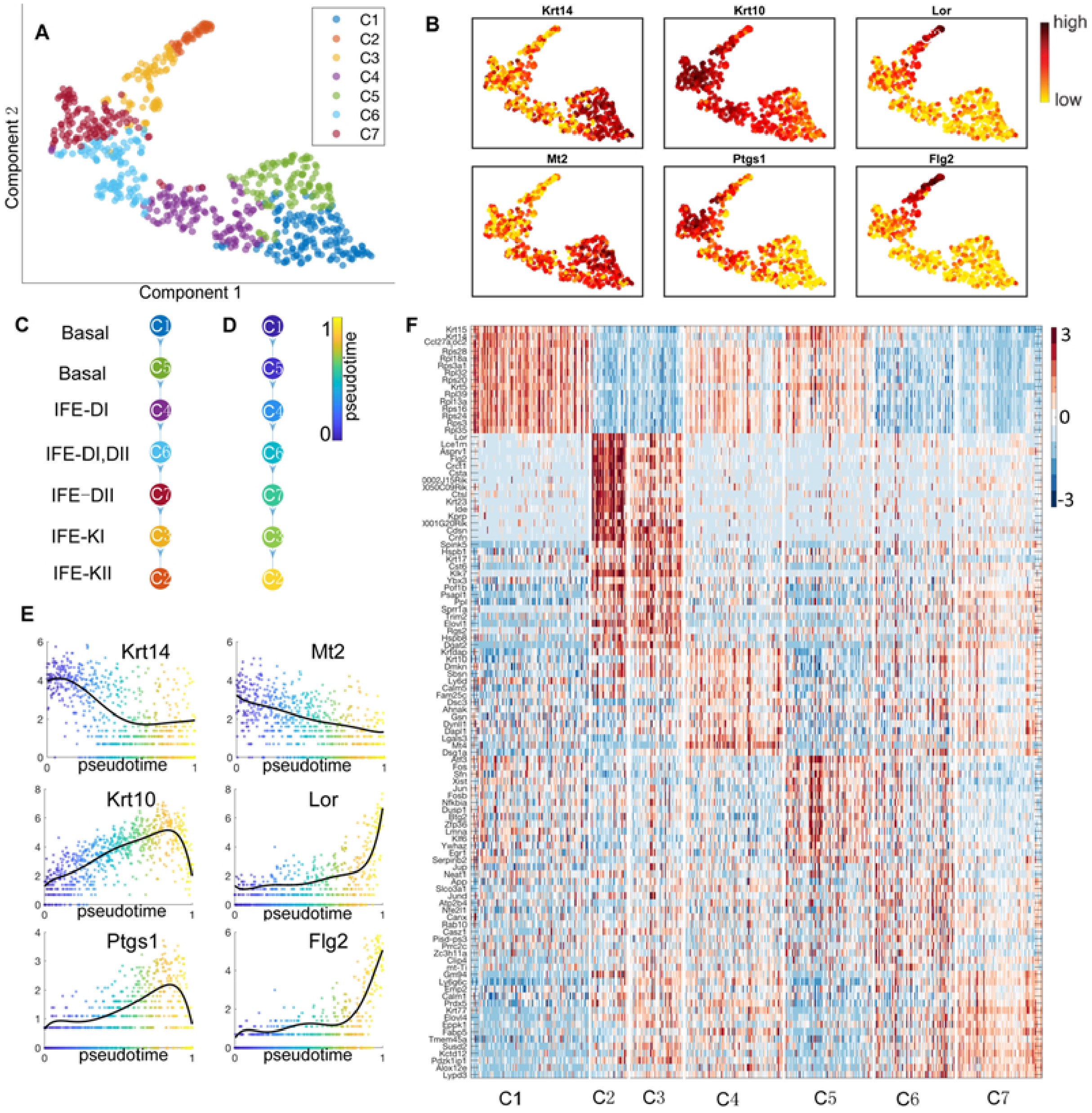

**Supplementary Figure S7.** Caption on next page.

**Supplementary Figure 7. Analysis of epidermal differentiation in the IFE.** [5] (A) Cell subpopulations inferred by SoptSC and the low dimensional visualization. (B) Selected marker genes and their expression in each population. (C) Lineage tree identified by SoptSC. (D) Pseudotime for cell states where the pseudotime of each subpopulation is calculated by the average of the temporal ordering of cells within the subpopulation. (E) Expression of selected marker genes along pseudotime. (F) Clustered gene-cell heatmap of genes from top 15 markers for each cluster identified by SoptSC.

#### 1.8 Supplementary Figure S8

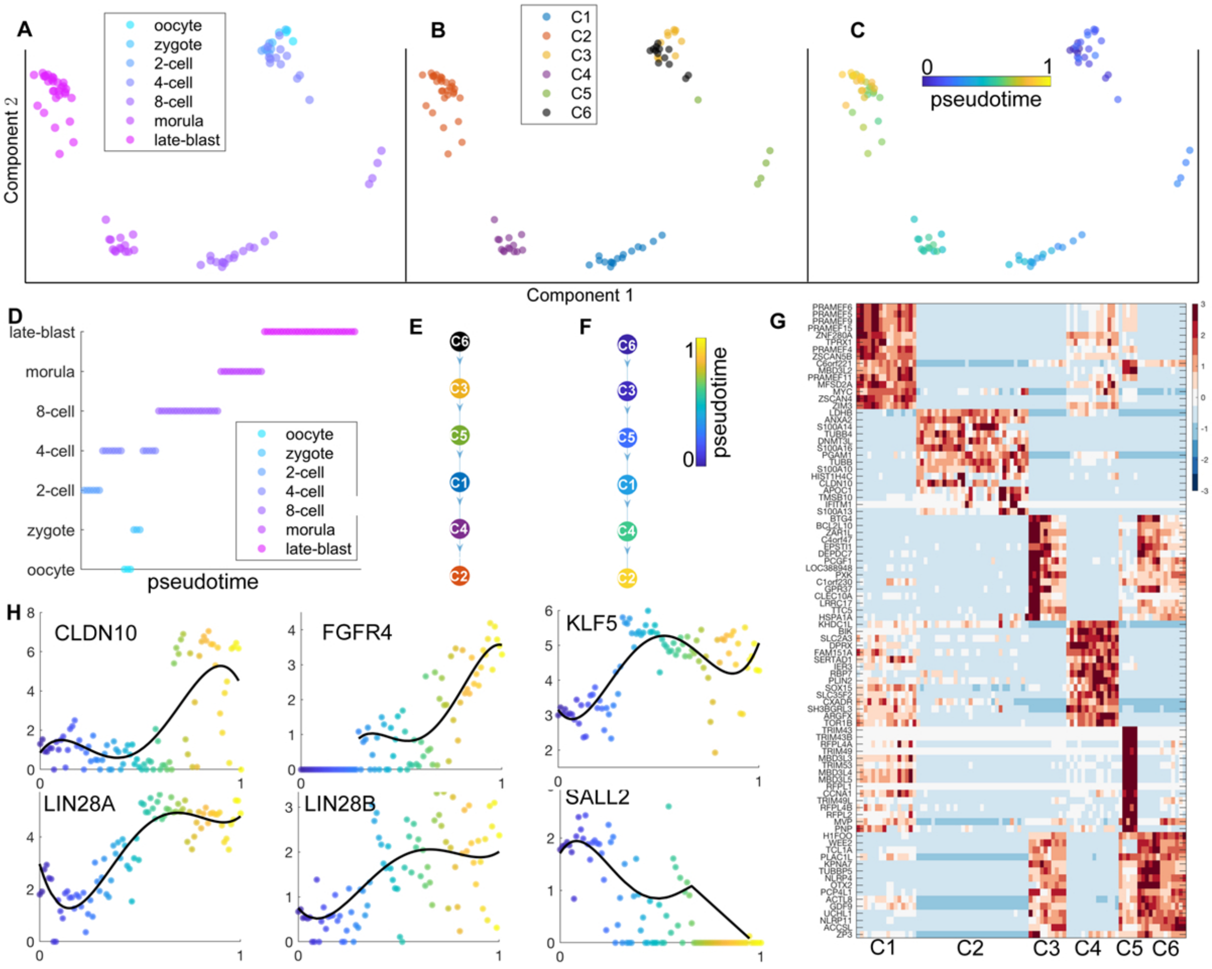

**Supplementary Figure S8. SoptSC identifies cell subpopulations, markers, lineage, and pseudotime in human early embryonic data.** [22] (A) Visualization of cell in two-dimensional space labeled by true experimental time information. (B) Unsupervised subpopulations identified by SoptSC. (C) Pseudotime inferred by SoptSC. (D) Pseudotemporal ordering of cells is compared with the known experimental stages. The Kendall rank correlation between the temporal ordering of cells inferred by SoptSC and the true experimental time is 0.84. (E) Lineage inferred by SoptSC, indicating a linear trajectory. (F) Pseudotime along lineage, where the pseudotime of each subpopulation is calculated by the average of the temporal ordering of cells within the subpopulation. (G) Clustered gene-cell heatmap of genes from top 15 markers for each cluster identified by SoptSC. (H) Expression of selected marker genes along pseudotime.

#### 1.9 Supplementary Figure S9

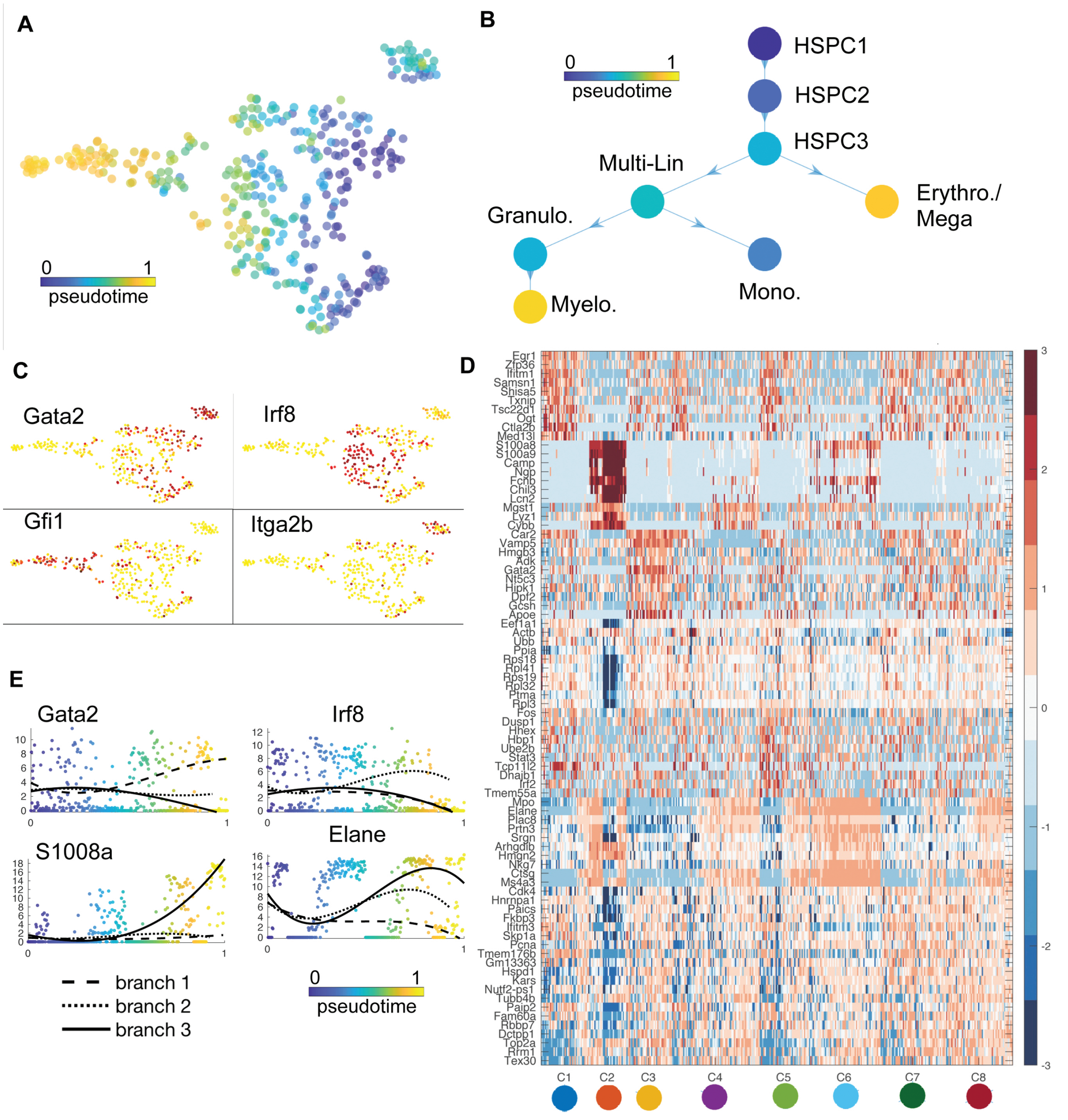

**Supplementary Figure S9.** Caption on next page.

**Supplementary Figure 9. Analysis of subpopulations, pseudotime, and lineage paths during myelopoiesis [15].** (A) Pseudotemporal ordering of hematopoietic cells by SoptSC. (B) Lineage hierarchy constructed by SoptSC. Colors correspond to the mean pseudotime value for the subpopulation. Subpopulation identities have been annotated according to marker gene expression. HSPC: hematopoietic stem/progenitor cells; Multi-Lin: mixed progenitor (see [15]); Mono: monocytic progenitor; Granulo: granulocytic progenitor; Myelo: myelocytic progenitor; Erythro: erythrocytic progenitor; Mega: megakaryocytic progenitor. (C) Gene expression of selected markers. (D) Clustered gene-cell heatmap of genes from top 15 markers for each cluster identified by SoptSC; (E) Expression of selected markers along pseudotime where branch 1 corresponds to HSPC1, HSPC2, HSPC3, Erythro./Mega; branch 2 corresponds to HSPC1, HSPC2, HSPC3, Multi-Lin, Mono.; and branch 3 corresponds to HSPC1, HSPC2, HSPC3, Granulo., Myelo.;

#### 1.10 Supplementary Figure S10

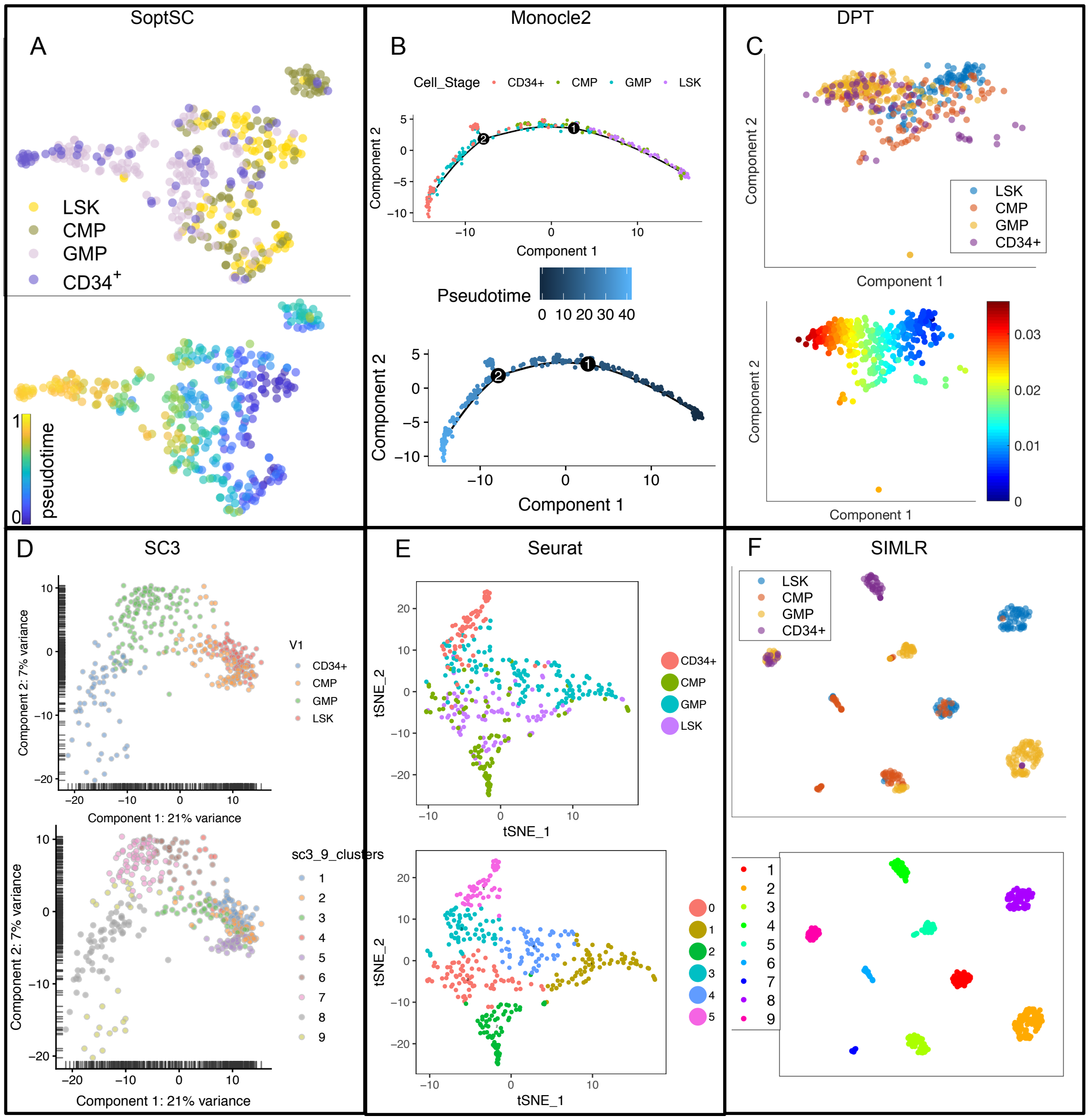

**Supplementary Figure S10. Pseudotime inference by SoptSC, Monocle2 and DPT; and clusters identified by SC3, Seurat, and SIMLR for Olsson et al. data [16].** (A, B, C) Visualization of two-dimensional projection of cells by (SoptSC, Monocle2, DPT) with true labels from the original study and pseudotime inferred by (SoptSC, Monocle2, DPT). (D, E, F) Visualization of low-dimensional projection of cells by (SC3,Seurat,SIMLR) with cell-stage labels and cluster labels identified by (SC3, Seurat, SIMLR).

##### 1.11 Supplementary Figure S11

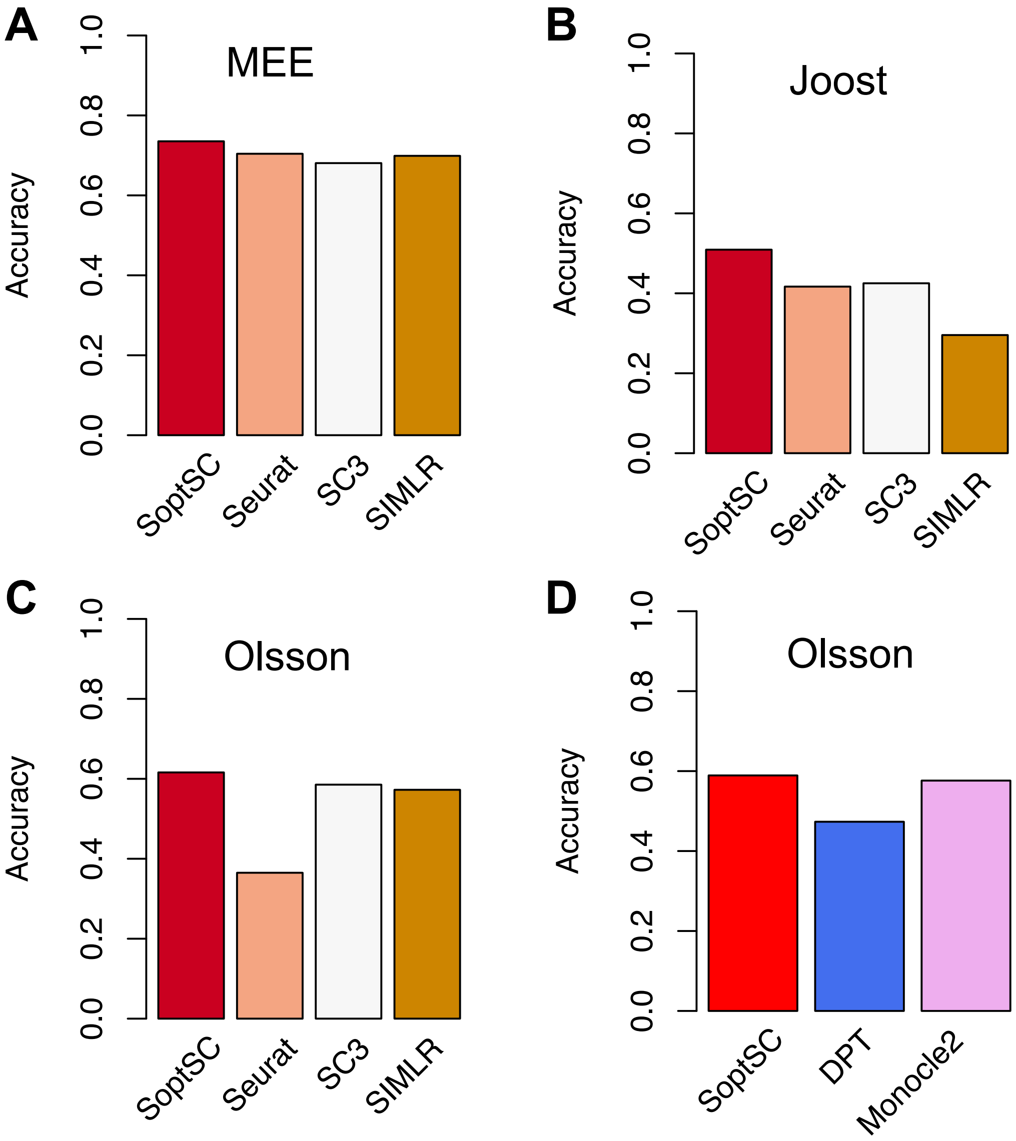

**Supplementary Figure S11. Comparison of performance of clustering for methods SoptSC, Seurat, SC3, SIMLR, and pseudotime inference for SoptSC, Monocle2, DPT** (A) Accuracy of clusters identified by SoptSC, Seurat, SC3 and SIMLR for mouse early embryonic data [4]. (B) Accuracy of clusters identified by SoptSC, Seurat, SC3 and SIMLR for IFE data [5]. (C) Accuracy of clusters identified by SoptSC, Seurat, SC3 and SIMLR for Olsson et al. data data [16]. (D) Accuracy of pseudotime inferred by SoptSC, Monocle2 and DPT for Olsson et al. data data [16].

#### 1.12 Supplementary Figure S12

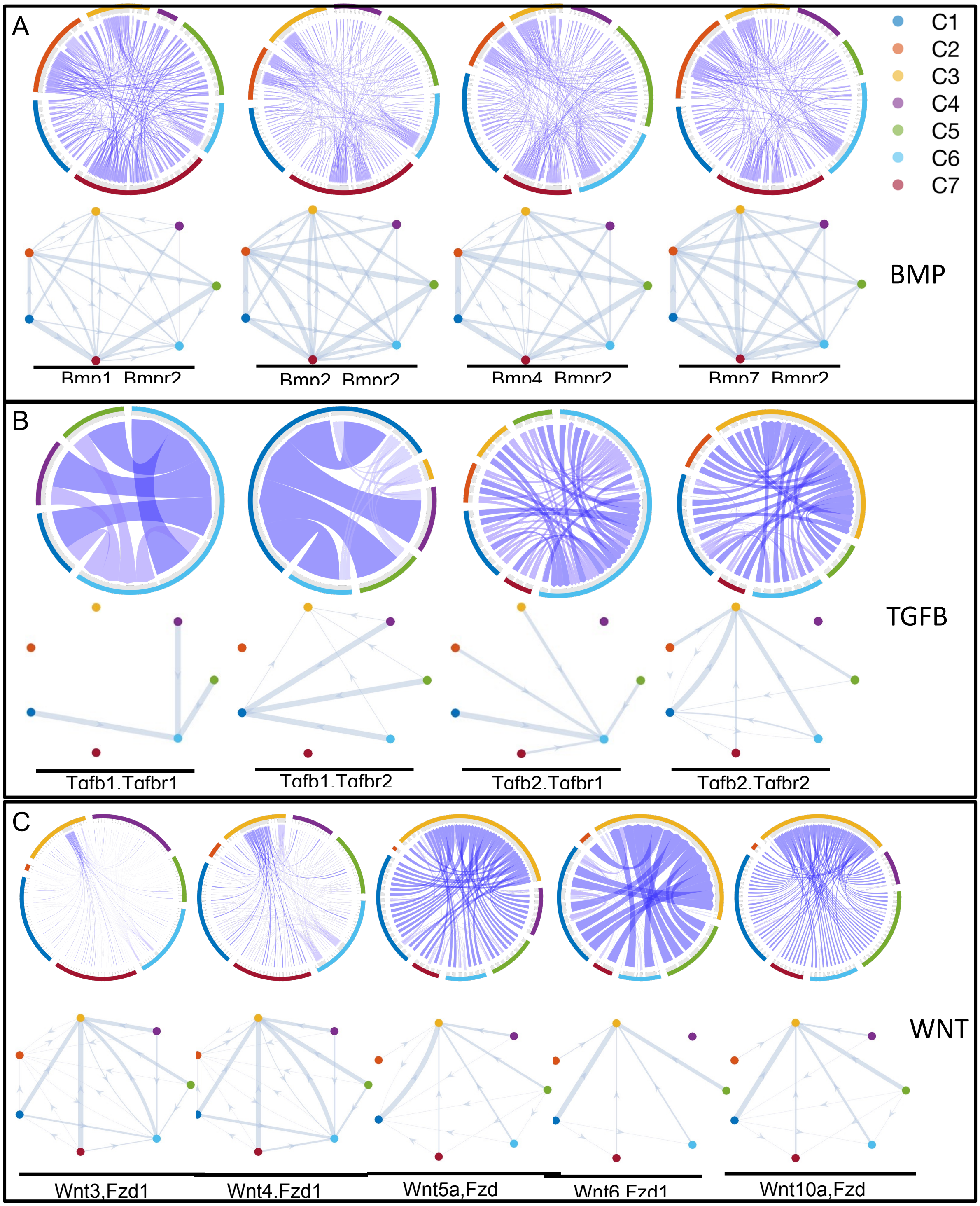

**Supplementary Figure S12. Single-cell signaling networks predicted for the BMP, TGF$\beta$, and Wnt pathways from Joost et al. data [5].** (A) Cell-cell signaling network and cluster-cluster signaling network for individual ligand-receptor pair from BMP pathway. Top row: single-cell signaling networks for ligand-receptor pairs, with edge weights corresponding to the probability of a signal passed between cells. Bottom row: cluster-to-cluster signaling interactions with edge weights corresponding to the probability of a signal passed between clusters. Colors correspond to the cluster labels. (B) Cell-cell signaling network and cluster-cluster signaling network for individual ligand-receptor pair from TGF$\beta$ pathway. (C) Cell-cell signaling network and cluster-cluster signaling network for individual ligand-receptor pair from Wnt pathway. Target gene list is summarized in **Table S2**.

#### 1.13 Supplementary Figure S13

**Bmp**

| | C1 | C2 | C3 | C4 | C5 | C6 | C7 |
|---|---|---|---|---|---|---|---|
| C1 | 1034 | 752 | 1833 | 2162 | 987 | 1880 | 3196 |
| C2 | 374 | 272 | 663 | 782 | 357 | 680 | 1156 |
| C3 | 682 | 496 | 1209 | 1426 | 651 | 1240 | 2108 |
| C4 | 594 | 432 | 1053 | 1242 | 567 | 1080 | 1836 |
| C5 | 1320 | 960 | 2340 | 2760 | 1260 | 2400 | 4080 |
| C6 | 638 | 464 | 1131 | 1334 | 609 | 1160 | 1972 |
| C7 | 748 | 544 | 1326 | 1564 | 714 | 1360 | 2312 |

**Tgfb**

| | C1 | C2 | C3 | C4 | C5 | C6 | C7 |
|---|---|---|---|---|---|---|---|
| C1 | 24 | 0 | 9 | 12 | 36 | 24 | 18 |
| C2 | 16 | 0 | 6 | 8 | 24 | 16 | 12 |
| C3 | 24 | 0 | 9 | 12 | 36 | 24 | 18 |
| C4 | 0 | 0 | 0 | 0 | 0 | 0 | 0 |
| C5 | 16 | 0 | 6 | 8 | 24 | 16 | 12 |
| C6 | 48 | 0 | 18 | 24 | 72 | 48 | 36 |
| C7 | 8 | 0 | 3 | 4 | 12 | 8 | 6 |

**Wnt**

| | C1 | C2 | C3 | C4 | C5 | C6 | C7 |
|---|---|---|---|---|---|---|---|
| C1 | 558 | 744 | 1302 | 372 | 186 | 1116 | 1488 |
| C2 | 78 | 104 | 182 | 52 | 26 | 156 | 208 |
| C3 | 225 | 300 | 525 | 150 | 75 | 450 | 600 |
| C4 | 561 | 748 | 1309 | 374 | 187 | 1122 | 1496 |
| C5 | 540 | 720 | 1260 | 360 | 180 | 1080 | 1440 |
| C6 | 492 | 656 | 1148 | 328 | 164 | 984 | 1312 |
| C7 | 951 | 1268 | 2219 | 634 | 317 | 1902 | 2536 |

**Supplementary Figure S13. Number of ligand-receptor pairs between clusters from Joost et al. data.** [5] Tables contain the number of ligand-receptor pairs between clusters identified by SoptSC for Bmp, Tgf-$\beta$ and Wnt pathways (members of which are summarized in **Table S2**).

##### 1.14 Supplementary Figure S14

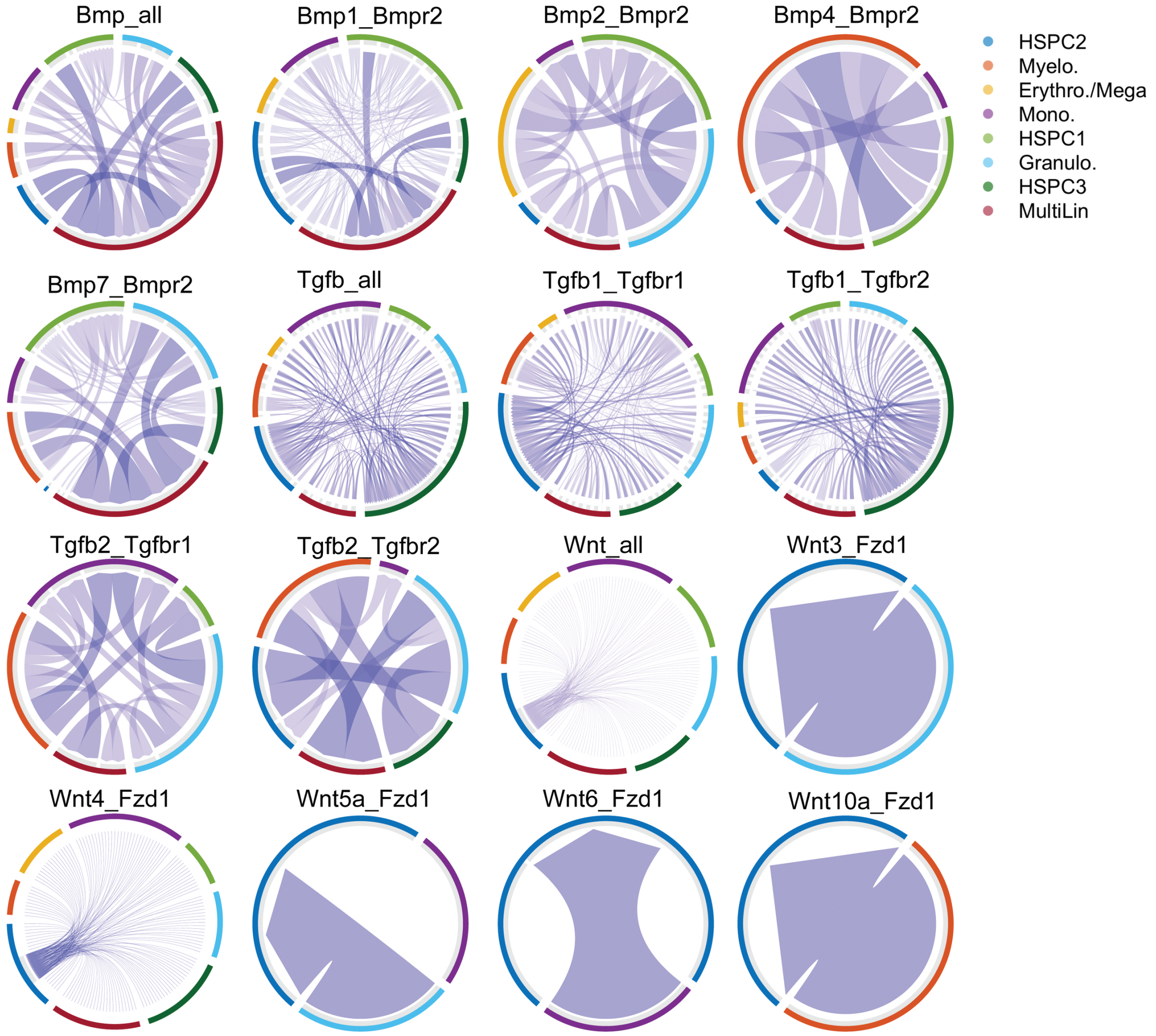

**Supplementary Figure S14. Cell-to-cell signaling networks predicted for pathways in data from Olsson et al. [15]** Single-cell signaling networks for ligand-receptor pairs, with edge weights corresponding to the probability of a signal passed between cells. Members of the pathways analyzed for Bmp, Tgf-$\beta$ and Wnt are summarized in **Table S3**.

#### 1.15 Supplementary Figure S15

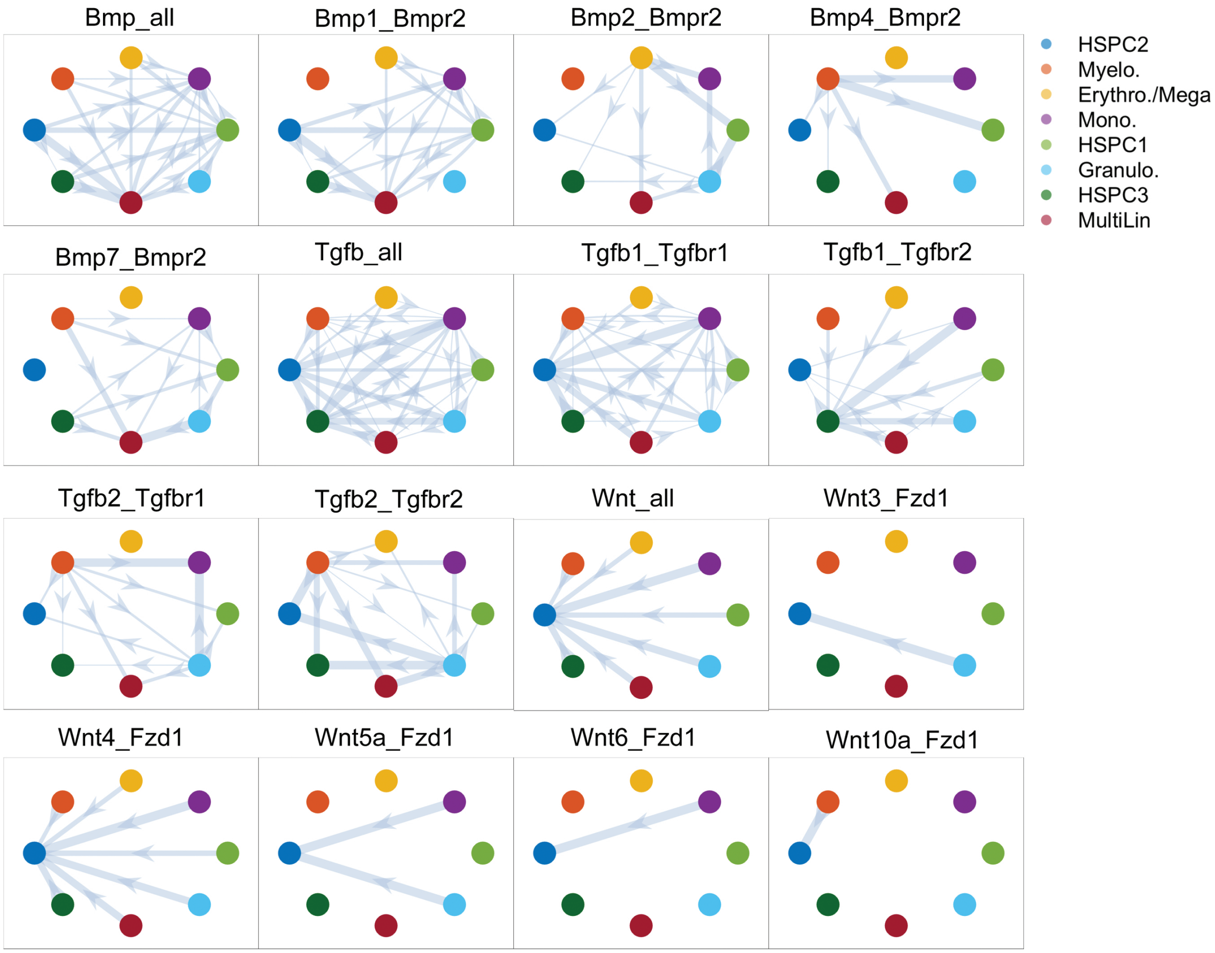

**Supplementary Figure S15. Cluster-to-cluster signaling networks predicted for pathways in data from Olsson et al.** [15] Bmp_all, Tgf$\beta$_all and Wnt_all represent the signaling network between clusters using all provided ligand-receptor pairs for the specific pathways.

#### 1.16 Supplementary Figure S16

| | C1 | C2 | C3 | C4 | C5 | C6 | C7 | C8 |
|---|---|---|---|---|---|---|---|---|
| C1 | 80 | 60 | 60 | 340 | 240 | 80 | 180 | 200 |
| C2 | 0 | 0 | 0 | 0 | 0 | 0 | 0 | 0 |
| C3 | 60 | 45 | 45 | 255 | 180 | 60 | 135 | 150 |
| C4 | 48 | 36 | 36 | 204 | 144 | 48 | 108 | 120 |
| C5 | 32 | 24 | 24 | 136 | 96 | 32 | 72 | 80 |
| C6 | 12 | 9 | 9 | 51 | 36 | 12 | 27 | 30 |
| C7 | 36 | 27 | 27 | 153 | 108 | 36 | 81 | 90 |
| C8 | 48 | 36 | 36 | 204 | 144 | 48 | 108 | 120 |

Bmp

| | C1 | C2 | C3 | C4 | C5 | C6 | C7 | C8 |
|---|---|---|---|---|---|---|---|---|
| C1 | 0 | 0 | 0 | 0 | 0 | 0 | 0 | 0 |
| C2 | 84 | 60 | 50 | 124 | 54 | 108 | 80 | 102 |
| C3 | 58 | 40 | 36 | 84 | 38 | 73 | 56 | 70 |
| C4 | 290 | 200 | 180 | 420 | 190 | 365 | 280 | 350 |
| C5 | 174 | 120 | 108 | 252 | 114 | 219 | 168 | 210 |
| C6 | 174 | 120 | 108 | 252 | 114 | 219 | 168 | 210 |
| C7 | 290 | 200 | 180 | 420 | 190 | 365 | 280 | 350 |
| C8 | 232 | 160 | 144 | 336 | 152 | 292 | 224 | 280 |

Tgfb

| | C1 | C2 | C3 | C4 | C5 | C6 | C7 | C8 |
|---|---|---|---|---|---|---|---|---|
| C1 | 108 | 0 | 0 | 0 | 0 | 108 | 0 | 0 |
| C2 | 52 | 0 | 0 | 0 | 0 | 52 | 0 | 0 |
| C3 | 88 | 0 | 0 | 0 | 0 | 88 | 0 | 0 |
| C4 | 174 | 0 | 0 | 0 | 0 | 174 | 0 | 0 |
| C5 | 104 | 0 | 0 | 0 | 0 | 104 | 0 | 0 |
| C6 | 105 | 0 | 0 | 0 | 0 | 105 | 0 | 0 |
| C7 | 124 | 0 | 0 | 0 | 0 | 124 | 0 | 0 |
| C8 | 120 | 0 | 0 | 0 | 0 | 120 | 0 | 0 |

Wnt

**Supplementary Figure S16. Number of ligand-receptor pairs between clusters from Olsson et al. data. [15]** Tables contain the number of ligand-receptor pairs between clusters identified by SoptSC for Bmp, Tgf-$\beta$ and Wnt pathways (members of which are summarized in **Table S3**. {C1,C2,C3,C4,C5,C6,C7,C8} corresponds to {HSPC2, Myelo., Erythro./Mega, Mono., HSPC1, Granulo., HSPC3, MultiLin }.

#### 1.17 Supplementary Figure S17

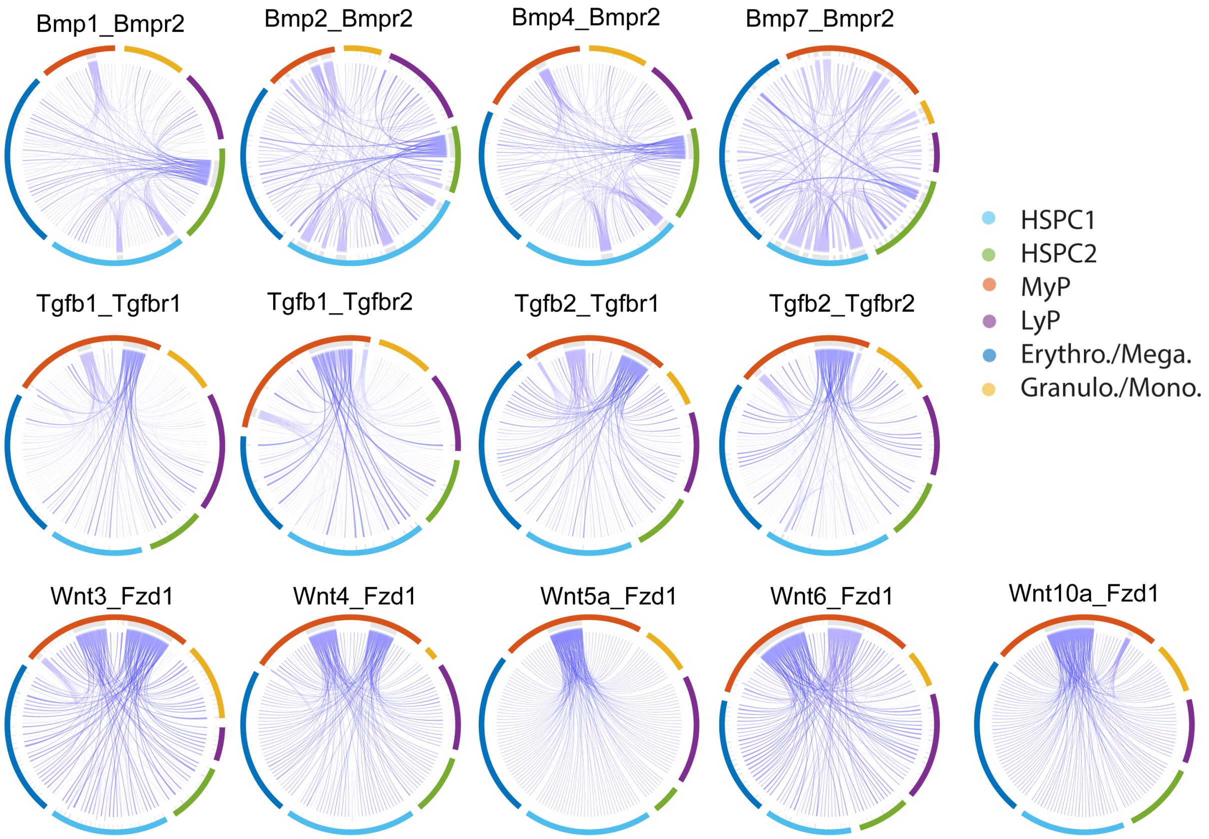

**Supplementary Figure S17. Cell-to-cell signaling networks predicted for pathways in data for mouse hematopoietic stem cell differentiation [14].** Single-cell signaling networks for ligand-receptor pairs, with edge weights corresponding to the probability of a signal passed between cells. Members of the pathways analyzed for Bmp, Tgf-$\beta$ and Wnt are summarized in **Table S4**.

#### 1.18 Supplementary Figure S18

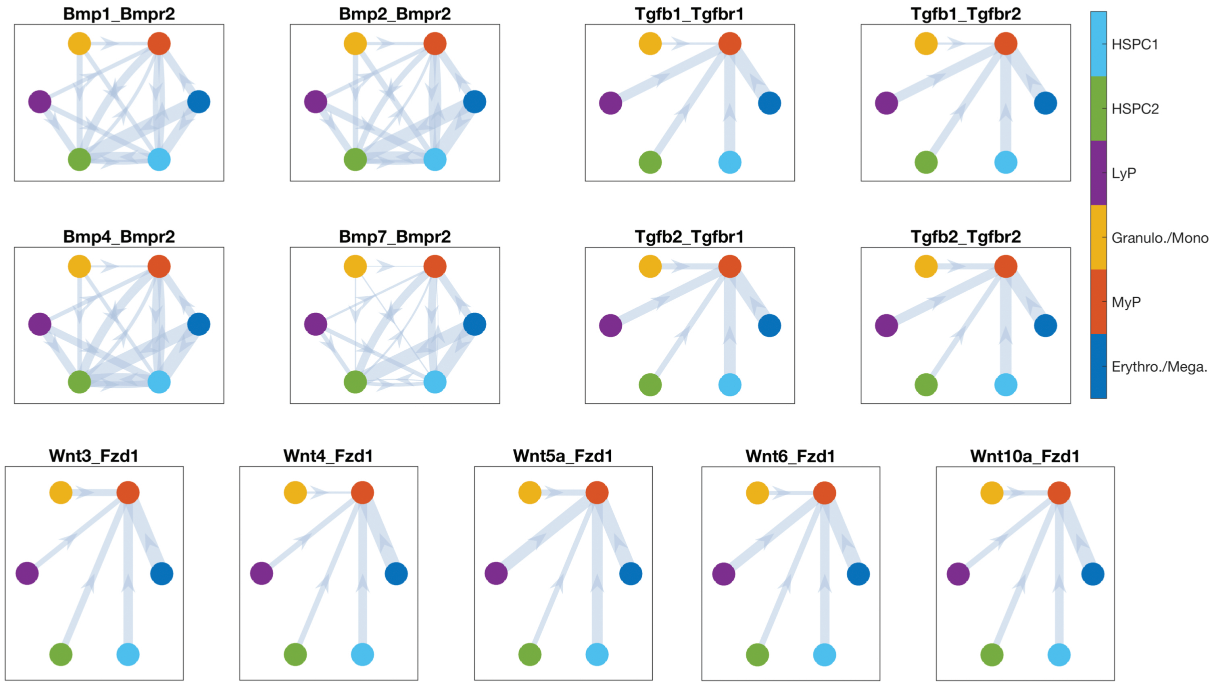

**Supplementary Figure S18. Cluster-to-cluster signaling networks predicted for pathways in data for mouse hematopoietic stem cell differentiation [14].** Bmp_all, Tgf$\beta$_all and Wnt_all represent the signaling network between clusters using all provided ligand-receptor pairs for the specific pathways.

#### 1.19 Supplementary Figure S19

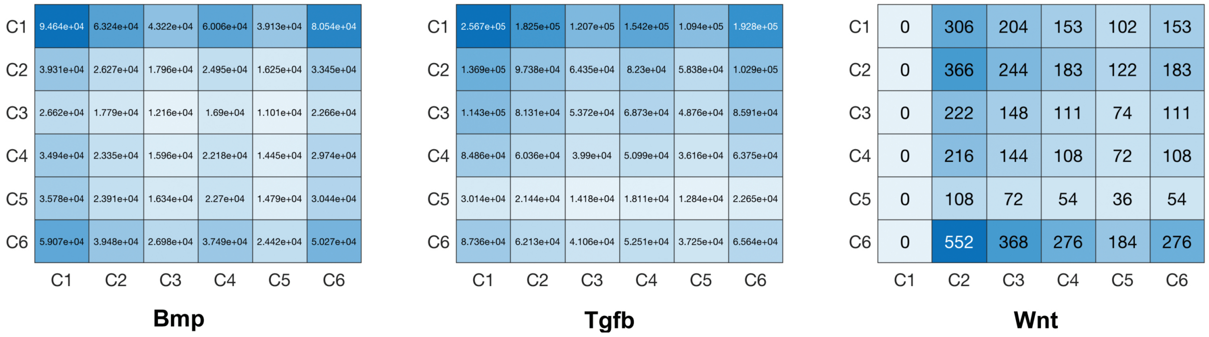

**Supplementary Figure S19. Number of ligand-receptor pairs between clusters for mouse hematopoietic stem cell differentiation [14].** Tables contain the number of ligand-receptor pairs between clusters identified by SoptSC for Bmp, Tgf-$\beta$ and Wnt pathways (members of which are summarized in **Table S4**. {C1,C2,C3,C4,C5,C6} corresponds to {Erythro./Mega., MyP, Granulo./Mono., LyP, HSPC2, , HSPC1}.

### 2 Supplementary Tables

#### 2.1 Supplementary Table S1

| Dataset | Number of Cells | Number of Genes | Number of Cell-types | Units |
|---|---|---|---|---|
| Yan [22] | 90 | 20214 | 7 | RPKM |
| Pollen [17] | 249 | 23730 | 4, 11 | TPM |
| Deng [2] | 268 | 22431 | 6, 10 | RPKM |
| Goolam [3] | 124 | 41480 | 5 | CPM |
| Kolodziejczyk [8] | 704 | 10685 | 3 | CPM |
| Treutlein [19] | 80 | 23271 | 5 | FPKM |
| Usoskin [20] | 622 | 25334 | 4, 8, 11 | RPM |
| Klein [7] | 2717 | 24175 | 4 | UMI |
| Zeisel [23] | 3005 | 19972 | 9 | UMI |

**Table S1. Summary of the characteristics of all the single cell datasets used in clustering performance comparison the paper (for Figure 2).**

#### 2.2 Supplementary Table S2

| Pathways | Ligand | Receptor | Target Genes |
|---|---|---|---|
| Tgf$\beta$ | Tgf$\beta$1 | Tgf$\beta$r1 | Zeb2, Smad2, Wnt4, Wnt11, Bmp7, Sox9, Notch1 |
| | Tgf$\beta$1 | Tgf$\beta$r2 | Zeb2, Smad2, Wnt4, Wnt11, Bmp7, Sox9, Notch1 |
| | Tgf$\beta$2 | Tgf$\beta$r1 | Zeb2, Smad2, Wnt4, Wnt11, Bmp7, Sox9, Notch1 |
| | Tgf$\beta$2 | Tgf$\beta$r2 | Zeb2, Smad2, Wnt4, Wnt11, Bmp7, Sox9, Notch1 |
| Bmp | Bmp1 | Bmpr2 | Crebbp, Fos, Id1, Jun, Runx1, Smad1, Smad5, Sox4, Cdh1 |
| | Bmp2 | Bmpr2 | Crebbp, Fos, Id1, Jun, Runx1, Smad1, Smad5, Sox4, Cdh1 |
| | Bmp4 | Bmpr2 | Crebbp, Fos, Id1, Jun, Runx1, Smad1, Smad5, Sox4, Cdh1 |
| | Bmp7 | Bmpr2 | SCrebbp, Fos, Id1, Jun, Runx1, Smad1, Smad5, Sox4, Cdh1 |
| Wnt | Wnt3 | Fzd1 | Ctnnb1, Lgr5, Runx2, Apc, Mmp7, Dkk1, Ccnd1 |
| | Wnt4 | Fzd1 | Ctnnb1, Lgr5, Runx2, Apc, Mmp7, Dkk1, Ccnd1 |
| | Wnt5a | Fzd1 | Ctnnb1, Lgr5, Runx2, Apc, Mmp7, Dkk1, Ccnd1 |
| | Wnt6 | Fzd1 | Ctnnb1, Lgr5, Runx2, Apc, Mmp7, Dkk1, Ccnd1 |
| | Wnt10a | Fzd1 | Ctnnb1, Lgr5, Runx2, Apc, Mmp7, Dkk1, Ccnd1 |

**Table S2. Signaling pathways used for generating cell-to-cell signaling networks and cluster-to-cluster signaling networks from Joost data [5].**

#### 2.3 Supplementary Table S3

| Pathways | Ligand | Receptor | Target Genes |
|---|---|---|---|
| Tgf$\beta$ | Tgf$\beta$1 | Tgf$\beta$r1 | Zeb2, Smad2, Wnt4, Wnt11, Bmp7, Sox9, Notch1 |
| | Tgf$\beta$1 | Tgf$\beta$r2 | Zeb2, Smad2, Wnt4, Wnt11, Bmp7, Sox9, Notch1 |
| | Tgf$\beta$2 | Tgf$\beta$r1 | Zeb2, Smad2, Wnt4, Wnt11, Bmp7, Sox9, Notch1 |
| | Tgf$\beta$2 | Tgf$\beta$r2 | Zeb2, Smad2, Wnt4, Wnt11, Bmp7, Sox9, Notch1 |
| Bmp | Bmp1 | Bmpr2 | Crebbp, Fos, Id1, Jun, Runx1, Smad1, Smad5, Sox4, Cdh1 |
| | Bmp2 | Bmpr2 | Crebbp, Fos, Id1, Jun, Runx1, Smad1, Smad5, Sox4, Cdh1 |
| | Bmp4 | Bmpr2 | Crebbp, Fos, Id1, Jun, Runx1, Smad1, Smad5, Sox4, Cdh1 |
| | Bmp7 | Bmpr2 | SCrebbp, Fos, Id1, Jun, Runx1, Smad1, Smad5, Sox4, Cdh1 |
| Wnt | Wnt3 | Fzd1 | Ctnnb1, Lgr5, Runx2, Apc, Mmp7, Dkk1, Ccnd1 |
| | Wnt4 | Fzd1 | Ctnnb1, Lgr5, Runx2, Apc, Mmp7, Dkk1, Ccnd1 |
| | Wnt5a | Fzd1 | Ctnnb1, Lgr5, Runx2, Apc, Mmp7, Dkk1, Ccnd1 |
| | Wnt6 | Fzd1 | Ctnnb1, Lgr5, Runx2, Apc, Mmp7, Dkk1, Ccnd1 |
| | Wnt10a | Fzd1 | Ctnnb1, Lgr5, Runx2, Apc, Mmp7, Dkk1, Ccnd1 |

**Table S3. Signaling pathways used for generating cell-to-cell signaling networks and cluster-to-cluster signaling networks from Olsson data [16].**

#### 2.4 Supplementary Table S4

| Pathways | Ligand | Receptor | Target Genes |
|---|---|---|---|
| Tgf$\beta$ | Tgf$\beta$1 | Tgf$\beta$r1 | Zeb2, Smad2, Wnt4, Wnt11, Bmp7, Sox9, Notch1 |
| | Tgf$\beta$1 | Tgf$\beta$r2 | Zeb2, Smad2, Wnt4, Wnt11, Bmp7, Sox9, Notch1 |
| | Tgf$\beta$2 | Tgf$\beta$r1 | Zeb2, Smad2, Wnt4, Wnt11, Bmp7, Sox9, Notch1 |
| | Tgf$\beta$2 | Tgf$\beta$r2 | Zeb2, Smad2, Wnt4, Wnt11, Bmp7, Sox9, Notch1 |
| Bmp | Bmp1 | Bmpr2 | Crebbp, Fos, Id1, Jun, Runx1, Smad1, Smad5, Cdh1 |
| | Bmp2 | Bmpr2 | Crebbp, Fos, Id1, Jun, Runx1, Smad1, Smad5, Cdh1 |
| | Bmp4 | Bmpr2 | Crebbp, Fos, Id1, Jun, Runx1, Smad1, Smad5, Cdh1 |
| | Bmp7 | Bmpr2 | SCrebbp, Fos, Id1, Jun, Runx1, Smad1, Smad5, Cdh1 |
| Wnt | Wnt3 | Fzd10 | Ctnnb1, Lgr5, Runx2, Apc, Mmp7, Dkk1, Ccnd1 |
| | Wnt4 | Fzd10 | Ctnnb1, Lgr5, Runx2, Apc, Mmp7, Dkk1, Ccnd1 |
| | Wnt5a | Fzd10 | Ctnnb1, Lgr5, Runx2, Apc, Mmp7, Dkk1, Ccnd1 |
| | Wnt8a | Fzd10 | Ctnnb1, Lgr5, Runx2, Apc, Mmp7, Dkk1, Ccnd1 |
| | Wnt8b | Fzd10 | Ctnnb1, Lgr5, Runx2, Apc, Mmp7, Dkk1, Ccnd1 |

**Table S4. Signaling pathways used for generating cell-to-cell signaling networks and cluster-to-cluster signaling networks for mouse hematopoietic stem cell differentiation [14].**

### 3 Extended Methods for SoptSC

Here we describe SoptSC, an optimization-based algorithm that enables the de novo detection of subpopulations, marker genes, cell lineage hierarchy, pseudotemporal ordering, and signaling pathways from single-cell gene expression datasets. SoptSC is based on the concept of similarity between cells, i.e. we find a low-rank representation of a cell (where we define a 'cell' here as the vector of gene expression values for a given cell) in relation to other cells within a neighborhood. The similarity score is thus defined by a set of linear coefficients (the solution of the low-rank optimization model) in a subspace given by its neighboring cells.

The SoptSC algorithm consists of several steps to perform clustering, lineage inference, pseudotemporal ordering and signaling network construction. All the steps work coherently with each other. In the first, a square matrix is constructed that describes the cell-to-cell similarities based on the input gene expression data. In the second step, low-rank approximations of the similarity matrix are calculated to define: ($i$) cell subpopulations within the data (rank-$k$, where $k$ is the number of subpopulations and identified in a unsupervised manner); ($ii$) ranked marker genes for each subpopulation. In the third step, cell-to-cell graph derived from the similarity matrix is designed to infer pseudotemporal ordering of cells. In the fourth step, cluster-to-cluster graph is constructed to identify the transition paths between cell subpopulations via minimal spanning tree (MST). At last, cell-to-cell as well as cluster-to-cluster signaling network is constructed for given ligand-receptor pairs via novel developed methods to compute cell-cell contact probability.

#### 3.1 Gene Filtering

SoptSC implements two steps to select highly variable genes. First, we remove genes that are expressed in less than $\alpha\%$ cells or expressed at least $1-\alpha\%$ cells ($\alpha = 6$ by default). This procedure is the same as SC3 for gene filter where the motivation is to reduce genes that are not informative for clustering. Second, we perform principal component analysis (PCA) on the single cell data with genes filtered in the first step. Genes with highest loadings in the first $k$ principal components are selected for downstream analysis. We set the value of $k$ as the index that the largest gap of the principal component variances occurs.

#### 3.2 Symmetric Non-negative Matrix Factorization (NMF) for Clustering

Let $S \in \mathbb{R}^{n\times n}$ be a non-negative symmetric matrix in which $S_{i,j}$ measures the similarity between cell $i$ and cell $j$. In order to classify cells into subpopulations based on their similarity, we use symmetric non-negative matrix factorization (NMF), which can be regarded as a graph-based clustering method and is widely used for data clustering [9, 10]. By applying NMF to $S$, the similarity matrix $S$ is decomposed into a product of a non-negative low rank matrix $H \in \mathbb{R}_{+}^{n\times k}$ and its transpose $H^{\top}$ via the optimization problem:

$$\begin{aligned} \mathcal{P}_2 : \min_{H\in\mathbb{R}^{n\times k}} \; & ||S - HH^{\top}||_F^2 \\ \text{s.t.} \quad & H \geq 0, \end{aligned}$$

where $k$ is the number of subpopulations of cells and $||\cdot||_F$ is the Frobenius norm. Such a NMF model is similar to spectral clustering, where the non-negative constraint here is replaced by an orthogonal constraint [9]. The low rank condition for $H$ is ideally suited for capturing the clustered nature of the cell subpopulations, i.e. by reordering $S$ according to the columns of $H$, a block-diagonal or near-block-diagonal structure can be obtained. We denote the reordered similarity matrix $S^B$. The structure of $S^B$ is such that cells within a block have high similarity to each other and low similarity to cells from other blocks. It can be shown that the solution of $\mathcal{P}_1$ is strictly block-diagonal when the data are clean and sampled from independent subspaces [11]. Due to this observation, the similarity matrix $S$ can be approximated by a sum of rank one matrices $H^i H^{i\top}, i = 1, 2, ..., k$, where $H = [H^1, H^2, ..., H^n]$, which can be obtained by solving the NMF problem $\mathcal{P}_2$ . Singular value decomposition is used to find $H_0$, an initial low-rank non-negative matrix required as an input for $\mathcal{P}_2$ [1]. If we now let $S = [S^1, S^2, ..., S^n]$ represent the columns of $S$, then the columns of $S$ can be approximated by the space spanned by the columns of $H$ as:

$$S^i \approx \sum_{j=1}^{k} H_{i,j} H^j.$$

Thus, the columns of $H$ represent a basis for $S$ in the (low rank) $k$-dimensional space, and the columns of $H^\top$ provide the coefficients for their corresponding columns of $S$ in the space spanned by the columns of $H$.

Since $H \geq 0$, each column of $H^\top$ can be viewed as a distribution for which the $i^{th}$ column $S^i$ has the component in the corresponding column of $H$. We can use $H^{\mathrm{T}}$ to classify the $N$ cells into $k$ subpopulations by assigning the $i^{th}$ cell to the $j^{th}$ subpopulation when the largest element among all components of the $i^{th}$ column of $H^{\mathrm{T}}$ lies in the $j^{th}$ position.

##### 3.3 Predicting the Number of Clusters from Data

Determining the number of clusters in a dataset is a fundamental problem extending far beyond the identification of cell populations from single cell data; many clustering algorithms still require the user to specify the number of clusters. We propose a method to automatically identify the number of clusters within a dataset based on properties of the graph Laplacian ($L$) and the consensus similarity matrix [21], which is similar to [12]. We consider a range of values: $k_i = \{k_1, k_2, ..., k_q\}$, which can be viewed as a prior distribution for the number of cell subpopulations.

It has been shown that the number of eigenvalues of $L$ equal to 0 is equivalent to the number of diagonal blocks of $L$ [21].

The steps required to determine the number of clusters $k$ are as follows:

1. Given the inputs $S$ and $k_i, i \in (1, 2, ..., q)$, partition the cells into $k_i$ subpopulations by solving the NMF problem $\mathcal{P}_2$ .

2. Find the consensus matrix [6, 13], $C$. For each $j \in (1, 2, ..., q)$, define a matrix $M^j$ by

$$M^j_{p,q} = \begin{cases} 1 & \text{if p and q belong to the same cluster} \\ 0 & \text{otherwise.} \end{cases}$$

The consensus matrix $C$ is then defined by

$$C = \sum_{j=1}^{q} M^j$$

3. Prune the consensus matrix as follows: set a tolerance $\tau \in [0, 0.5]$, and let $C_{i,j} = 0$ if $C_{i,j} \leq \tau q$. This increases the robustness of consensus clustering to biological noise.

4. Compute the graph Laplacian $L$ and its eigenvalues, given the identity matrix $I$ and a diagonal matrix $D$ such that:

$$L = I - D^{-1/2} C D^{-1/2}$$

$$\text{with } D_{ii} = \sum_{j=1}^{n} C_{i,j}.$$

5. Find ($i$) the number of eigenvalues that are close to zero, and ($ii$) the index at which the largest eigenvalue gap occurs [21].

An initial estimate for $k$ is given by ($i$). In cases where there may be significant sources of noise in the data, or where other uncertainties exist, we can use ($ii$) instead as an estimate of $k$. Especially for cases displaying a prominent largest eigenvalue gap (see Supplementary Figure S1), ($ii$) can provide a better estimate of the subpopulation structure present in the data. For all analyses performed below, we choose a prior $k_i$ so the number of clusters ranges from 1 to 25, and we set the tolerance $\tau = 0.3$.

### 4 Extended Details on Data Analysis

#### 4.1 Details of Data Analysis by SoptSC

All the results in this paper is run under MATLAB R2017b on Mac Pro (Late 2013) with 3.5 GHz 6-Core Intel Xeon E5.

For all the datasets [2, 3, 7, 8, 17, 19, 20, 22, 23] used in evaluating the performance clustering (**Fig. 2**), we set the parameter value for gene selection as $\alpha = 0$ (for datasets from Treutlein[19] and Yan[22]) and $\alpha = 1$ for all the other datasets. In all cases, the number of selected genes is set as 2000.

For single cell qPCR data from mouse early embryo (e.g., [4]), we remove the first two control genes (actb, ahcy) and use the data with the rest of 46 genes as a input for SoptSC.

For scRNA-seq data from human early embryo (ref. [22]), we selected genes that expressed at least 6 cells and at most among the overall 88 cells, which induces 11517 genes to be used in the downstream analysis.

For Joost data set (ref. [5]), we selected 3000 genes for downstream analysis based on our gene filtering technique with parameter $\alpha$ being set as $0.03 \times N$ where $N$ represents the number of cells in the data.

For the Olsson data set (ref. [16]), we selected 2000 genes for downstream analysis based on our gene filtering technique with $\alpha = 0$.

For scRNA-seq data from Nesterowa (ref. [14]), we selected 3000 genes for downstream analysis based on our gene filtering techniques with $\alpha = 0.03 \times N$ where $N$ is the number of cells.

#### 4.2 Details of Data Analysis by SC3

SC3 is run in R under version 3.4.3. For all the datasets used in evaluating the performance clustering in the paper, we used the default setting.

#### 4.3 Details of Data Analysis by Seurat

For all datasets used to evaluate the performance of clustering against different methods (Figure 2), we implement the parameters setting for Seurat as follows. To initialize the Seurat object with the raw (non-normalized data), we keep all genes expressed in at least 3 cells and keep all cells with at least 200 detected genes To select highly variable genes for initial clustering of cells, we performed Principal Component Analysis (PCA) on the scaled data for all genes included in the previous step. We set x.low.cutoff = 0, y.cutoff = 0.8 in FindVariableGenes function. For clustering, we used the function FindClusters using 10 PCs with resolution 0.8. Nonlinear dimensionality reduction method, namely tSNE, was applied to the scaled matrix for visualization of cells in two-dimensional space using first 10 PC components.

For mouse early embryonic data [4], Joost et al. data [5] and Olsson et al. data [15] we used 3 PCs with resolution 0.8.

##### 4.4 Details of Data Analysis by SIMLR

We run SIMLR in MATLAB with default setting for all datasets used in the paper.

##### 4.5 Detailed Data Analysis by Monocle2

For two embryonic datasets used in pseudotime comparison, we run Monocle2 for the reduction method set as "DDRTree" with parameters pseudo expr_set as 0 and max_components set as 2. For Joost et al. [5] and Olsson et al. [15] datasets, we selected ordering genes with parameters mean_expression greater than 0.2 and dispersion_empirical larger than 0.5 ∗ dispersion_fit For Shalek et al. [18] data, we selected ordering genes with parameters mean_expression greater than 1 and dispersion_empirical larger than 1.5 ∗ dispersion_fit

##### 4.6 Details of Data Analysis by DPT

We run DPT in MATLAB with default setting for all datasets used in the paper.

15. Andre Olsson, Meenakshi Venkatasubramanian, Viren K Chaudhri, Bruce J Aronow, Nathan Salomonis, Harinder Singh, and H Leighton Grimes. Single-cell analysis of mixed-lineage states leading to a binary cell fate choice. *Nature*, 2016.

16. Andre Olsson, Meenakshi Venkatasubramanian, Viren K Chaudhri, Bruce J Aronow, Nathan Salomonis, Harinder Singh, and H Leighton Grimes. Single-cell analysis of mixed-lineage states leading to a binary cell fate choice. *Nature*, 537(7622):698–702, August 2016.

17. Alex A Pollen, Tomasz J Nowakowski, Joe Shuga, Xiaohui Wang, Anne A Leyrat, Jan H Lui, Nianzhen Li, Lukasz Szpankowski, Brian Fowler, Peilin Chen, et al. Low-coverage single-cell mrna sequencing reveals cellular heterogeneity and activated signaling pathways in developing cerebral cortex. *Nature biotechnology*, 32(10):1053, 2014.

18. Alex K Shalek, Rahul Satija, Joe Shuga, John J Trombetta, Dave Gennert, Diana Lu, Peilin Chen, Rona S Gertner, Jellert T Gaublomme, Nir Yosef, Schraga Schwartz, Brian Fowler, Suzanne Weaver, Jing Wang, Xiaohui Wang, Ruihua Ding, Raktima Raychowdhury, Nir Friedman, Nir Hacohen, Hongkun Park, Andrew P May, and Aviv Regev. Single-cell RNA-seq reveals dynamic paracrine control of cellular variation. *Nature*, June 2014.

19. Barbara Treutlein, Doug G Brownfield, Angela R Wu, Norma F Neff, Gary L Mantalas, F Hernan Espinoza, Tushar J Desai, Mark A Krasnow, and Stephen R Quake. Reconstructing lineage hierarchies of the distal lung epithelium using single-cell rna-seq. *Nature*, 509(7500):371, 2014.

20. Dmitry Usoskin, Alessandro Furlan, Saiful Islam, Hind Abdo, Peter Lönnerberg, Daohua Lou, Jens Hjerling-Leffler, Jesper Haeggström, Olga Kharchenko, Peter V Kharchenko, et al. Unbiased classification of sensory neuron types by large-scale single-cell rna sequencing. *Nature neuroscience*, 18(1):145, 2015.

21. Ulrike Von Luxburg. A tutorial on spectral clustering. *Statistics and computing*, 17(4):395–416, 2007.

22. Liying Yan, Mingyu Yang, Hongshan Guo, Lu Yang, Jun Wu, Rong Li, Ping Liu, Ying Lian, Xiaoying Zheng, Jie Yan, et al. Single-cell rna-seq profiling of human preimplanta-

tion embryos and embryonic stem cells. *Nature structural & molecular biology*, 20(9):1131–1139, 2013.

23. Amit Zeisel, Ana B. Muñoz-Manchado, Simone Codeluppi, Peter Lönnerberg, Gioele La Manno, Anna Juréus, Sueli Marques, Hermany Munguba, Liqun He, Christer Betsholtz, Charlotte Rolny, Gonçalo Castelo-Branco, Jens Hjerling-Leffler, and Sten Linnarsson. Cell types in the mouse cortex and hippocampus revealed by single-cell rna-seq. *Science*, 347(6226):1138–1142, 2015.
